# MRP5 and MRP9 Play a Concerted Role in Male Reproduction and Mitochondrial Function

**DOI:** 10.1101/2021.06.19.449033

**Authors:** Ian Chambers, Praveen Kumar, Jens Lichtenberg, Pengcheng Wang, Jianshi Yu, John Phillips, Maureen Kane, David Bodine, Iqbal Hamza

**Author notes:** **Corresponding Author:** Iqbal Hamza, **Email:**. **Author Contributions:** IGC: Conceptualization, Methodology, Investigation, Formal analysis, Validation, Visualization, Writing—original draft, Writing—review and editing PK: Conceptualization, Methodology, Investigation, Formal analysis, Validation JL: Methodology, Investigation, Formal analysis, Validation, Visualization PW: Methodology, Investigation, Formal analysis JY: Methodology, Investigation, Formal analysis JP: Conceptualization, Methodology, Investigation, Formal analysis, Validation, Resources, Funding acquisition, Writing—review and editing MAK: Conceptualization, Methodology, Investigation, Formal analysis, Validation, Resources, Funding acquisition, Writing—review and editing DB: Conceptualization, Methodology, Investigation, Formal analysis, Validation, Resources, Funding acquisition, Writing—review and editing IH: Conceptualization, Methodology, Investigation, Formal analysis, Validation, Visualization, Resources, Software, Supervision, Funding acquisition, Writing—original draft, Writing—review and editing.

## Abstract

Multidrug Resistance Proteins (MRPs) are transporters that play critical roles in cancer even though the physiological substrates of these enigmatic transporters are poorly elucidated. In *Caenorhabditis elegans*, MRP5/ABCC5 is an essential heme exporter as *mrp-5* mutants are unviable due to their inability to export heme from the intestine to extra-intestinal tissues. Heme supplementation restores viability of these mutants but fails to restore male reproductive deficits. Correspondingly, cell biological studies show that MRP5 regulates heme levels in the mammalian secretory pathway even though MRP5 knockout (KO) mice do not show reproductive phenotypes. The closest homolog of MRP5 is MRP9/ABCC12, which is absent in *C. elegans* raising the possibility that MRP9 may genetically compensate for MRP5. Here, we show that MRP5 and MRP9 double KO mice are viable but reveal significant male reproductive deficits. Although MRP9 is highly expressed in sperm, MRP9 KO mice show reproductive phenotypes only when MRP5 is absent. Both ABCC transporters localize to mitochondrial-associated membranes (MAMs), dynamic scaffolds that associate the mitochondria and endoplasmic reticulum. Consequently, DKO mice reveal abnormal sperm mitochondria with reduced mitochondrial membrane potential and fertilization rates. Metabolomics show striking differences in metabolite profiles in the DKO testes and RNA-seq show significant alterations in genes related to mitochondrial function and retinoic acid metabolism. Targeted functional metabolomics reveal lower retinoic acid levels in the DKO testes and higher levels of triglycerides in the mitochondria. These findings establish a model in which MRP5 and MRP9 play a concerted role in regulating male reproductive functions and mitochondrial sufficiency.

**Significance Statement:** MRPs are typically implicated in cancer biology. Here, we show that MRP9 and MRP5 localize to mitochondrial-associated membranes and play a concerted role in maintaining mitochondrial homeostasis and male reproductive fitness. Our work fills in significant gaps in our understanding of MRP9 and MRR5 with wider implications in male fertility. It is plausible that variants in these transporters are associated with male reproductive dysfunction.

## Introduction

Multidrug Resistance-associated Proteins (MRPs) belong to the type C subclass of the ATP Binding Cassette (ABCC) transporters, a highly conserved family of proteins that have been studied for decades for their role in cancers (1–7). MRPs are notoriously upregulated in a number of malignancies because of their ability to efflux a wide range of synthetic and chemotherapeutic agents, conferring cancer cells resistance to drug treatments (3, 8–11). Due to their critical importance in human health and clinical outcomes, myriads of MRP substrates have now been identified, particularly from *in vitro* studies of tumor cell lines (3, 4, 6, 12, 13). However, the promiscuity of drugs effluxed in this context is likely a functional consequence of cancer cells hijacking these active transporters rather than that being their true evolutionary role. Therefore, a major outstanding item remaining in the MRP field is the determination of true physiological functions and substrates for a number of these transporters.

Previous work from our group shed some light on the function of one such ABCC transporter, MRP5/ABCC5, where studies in worms, yeast, zebrafish and mammalian cells demonstrated it to be a heme exporter (14). Prior to this discovery, the role of MRP5 appeared to be that of cyclic nucleotide export from cells as demonstrated by transport of cGMP in membrane vesicles expressing human MRP5 *in vitro* (12, 15). However, the generation of MRP5 null mice demonstrated little to no contribution in cGMP efflux and no observable overt phenotypes, leaving the physiological relevance of this transporter unclear outside its *in vitro* or cancer cell context (16, 17). This is not the case in *C. elegans*, a heme auxotroph, where MRP-5 is absolutely essential for intercellular heme trafficking and knockout of *mrp*-5 results in heme deficiency and embryonic lethality (14).

While cell biological studies in mouse embryonic fibroblasts from MRP5 KO mice confirmed that MRP5 indeed plays a conserved role in heme export i.e. transporting heme into the secretory pathway for incorporation into hemoproteins or exporting it from the cell across the plasma membrane (14, 18–20), MRP5 KO mice do not show any heme-related phenotypes *in vivo*. A postulated hypothesis to explain these differences could be genetic compensation by other closely related transporters, which are phylogenetically conserved in higher order eukaryotes but absent in the *C. elegans* genome (14, 21). A candidate for this proposed compensation and further functional delineation is MRP9/ABCC12, the closest relative of MRP5 and one of the last of the ABCC proteins to have been discovered (6, 22, 23). MRP9 is highly expressed in male reproductive tissues but fails to effectively efflux drugs *in vitro*, and its endogenous function remains to be elucidated (6, 24, 25).

In this work, we generate a MRP9 knockout and MRP9 / MRP5 double knockout (DKO) mouse models to fully characterize these transporters’ physiological functions. Loss of MRP9 and MRP5 disrupts mitochondrial homeostasis and regulatory pathways in the testes, with RNA-seq and metabolomic analyses revealing large-scale perturbation in genes and metabolism related to mitochondrial function. Together, these results establish a concerted and critical role for MRP9 and MRP5 in male reproduction and mitochondrial metabolism, implying that the ancestral role of MRP5 in *C. elegans* reproduction was co-opted in the vertebrate ancestor.

## Results

### Loss of MRP9 in mice induces the expression of MRP5 in the testes

*C. elegans mrp-5(ok2067)* deletion mutants are lethal unless rescued with dietary heme supplementation (14). Serendipitously, we found that even in the presence of heme *mrp-5(ok2067)* male worms are defective in mating **(Figures S1A and S1B)**. These worms exhibit defective mating apparatus with stunted tail fan formation, reduced numbers of rays, and spicules which fail to retract despite normal mating behaviors. Given that hermaphrodite *mrp-5* knockout worms mate normally with wildtype males (not shown), it would appear the defect is paternal. For these reasons, we hypothesized that MRP-5 may have additional functions related to male reproduction outside of its traditional role in heme transport and homeostasis. MRP5 knockout mice however show no reproductive deficiencies, prompting inquiry into other ABCC transporters that may genetically compensate for the loss of MRP5 in higher order vertebrates.

Phylogenetic analysis of ABCC transporters across metazoans confirmed that members of the short MRP clade (ABCCs - 4, 5, 7, 11, 12) cluster tightly together across species indicating their evolutionary relation (6, 14). Within this subclade, *C. elegans* has only MRP-5 whereas most vertebrates have an additional paralog, MRP9 **(Figure 1A)**, which is conserved in all higher-order vertebrates but remains to be fully characterized *in vivo.* We therefore generated MRP9 mutant mice via CRISPR/Cas9 to explore MRP9’s endogenous functions. Utilizing guide RNAs targeted to the second exon of *Abcc12* we were able to generate 11 founder animals with various indels at the target site and a 17 bp insertion line was selected for further characterization **(Figures 1B and S1C)**. RT-PCR confirmed that the frameshift-inducing mutation was transcribed with no alternate start sites or splice variants identified **(Figure S1D)**.

**Figure 1.**
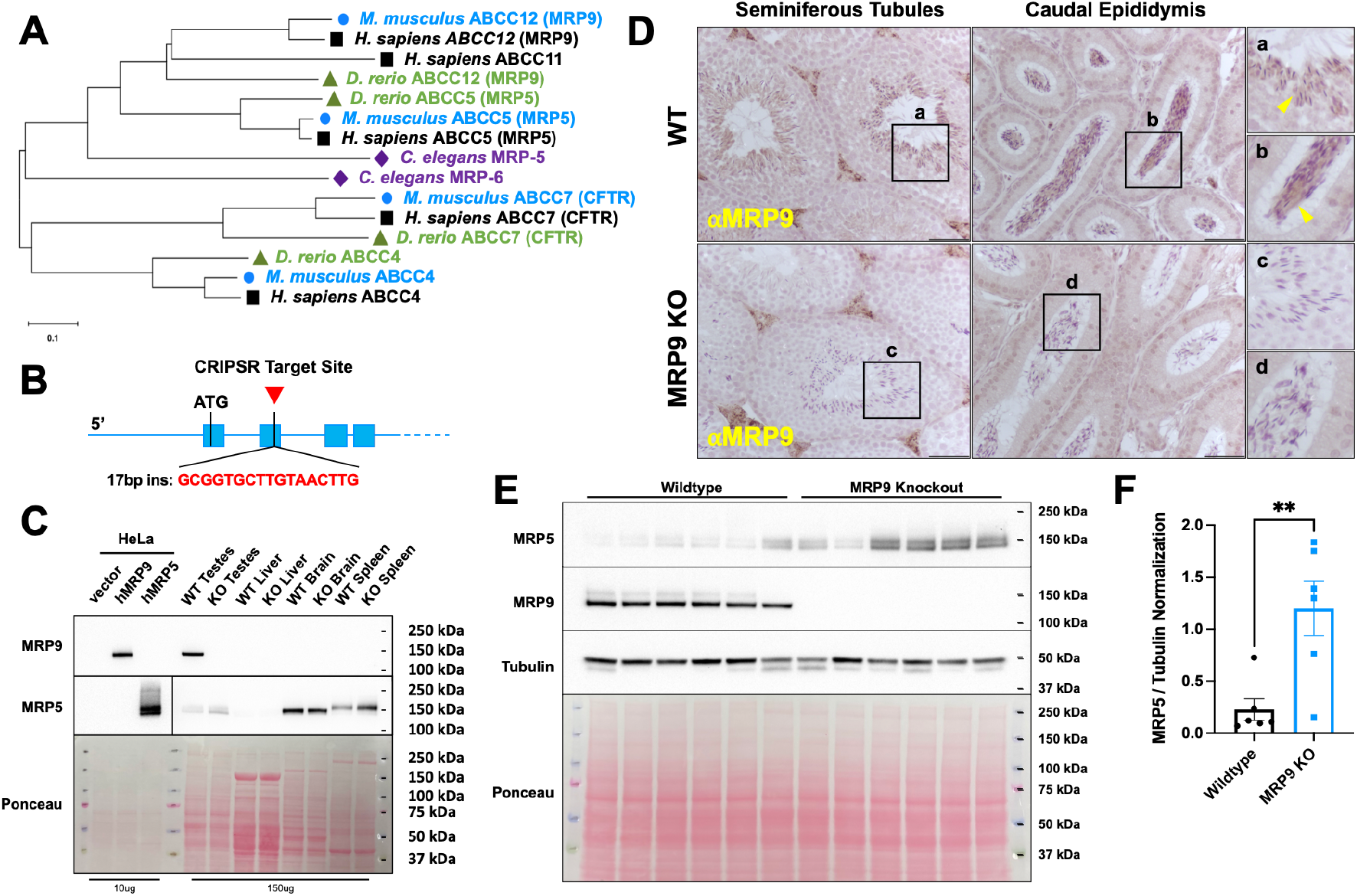
Ablation of MRP9 in mice induces MRP5 expression in the testes. **{A}** Amino acid FASTA sequences for MRPs in humans, mice, zebrafish and worms were aligned and used to generate the optimal phylogenetic tree shown. The evolutionary history was inferred using the Neighbor-Joining method. The tree is drawn to scale, with branch lengths in the same units as those of the evolutionary distances used to infer the phylogenetic tree. The evolutionary distances were computed using the Poisson correction method and are in the units of the number of amino acid substitutions per site. **{B}** Gene loci schematic of 5’ UTR and the first four exons of *Abcc12* indicating the CRISPR target site in second exon for the generation of knockout MRP9 mice. **{C}** Immunoblot of protein lysates from transfected HeLa cells and Wildtype and MRP9 KO tissues probed for MRP9 and MRP5. **{D}** MRP9 immunohistochemistry analysis of paraffin-embedded testes tissue sections from WT and MRP9 KO mice. Tissue sections were probed for MRP9 and lightly counterstained with hematoxylin. Images are representative of at least 3 mice; scale bar equals 100 μm. **{E}** Immunoblot of protein lysates from wildtype and MRP9 KO testes (n=6) probed for MRP5, MRP9 and Tubulin. **{F}** Quantification of MRP5 protein expression in wildtype and MRP9 KO testes from subpanel E, normalized to Tubulin; *P* value = 0.0061.

*Abcc12−/−* animals are viable and reach adulthood with no overt phenotypes. Immunoblots of lysates from testes of adult mice confirmed no detectable MRP9 protein in knockout animals **(Figure 1C)**. Immunohistochemistry of tissue sections showed MRP9 in both the maturing and developed spermatozoa of seminiferous tubules and caudal epididymis, respectively, consistent with the findings reported by Ono et al **(Figure 1D)** (24). Recent single cell RNAseq datasets of the testis niche further confirmed these findings as the gene expression profile of *Abcc12* is distinctly germ cell specific **(Figure S1E)** (26). This is in contrast to *Abcc5*, which appears to be normally expressed in early spermatogonia precursors as well as Sertoli and Leydig cells **(Figure S1E)**. Immunoblotting of lysates from additional tissues of wildtype and KO mice revealed the testes as the only organ with any appreciable amounts of both, MRP9 and MRP5 **(Figure 1C)**. Importantly, MRP9 KO testes showed elevated levels of MRP5 raising the notion that MRP5 could be compensating for MRP9 deficiency in the testes **(Figures 1E and 1F)**.

### MRP9 and MRP5 are required for male reproductive fitness

MRP9 KO mice do not show any statistically significant defects in reproduction, with expected litter sizes and normal Mendelian F1 segregation when intercrossed [n = 166, Chi squared = 0.976, *P* value = 0.614] **(Figure 2A)**. MRP9 KO crossed to MRP5 KO mice successfully generated DKO mice which reached adulthood with no overt phenotypes. Although DKO mice were viable, they were deficient in their reproductive capacity; the average number of pups from intercrosses of DKO mice were significantly reduced compared to WT [n > 33 litters, *P* value < 0.0001] **(Figure 2B)**. Interestingly, fecundity was restored back to single KO levels when DKO females were crossed with WT or double heterozygous males alone, suggestive that the failure may be paternal [n > 21 litters, *P* value < 0.0001] **(Figure 2B)**. Furthermore, DKO male mice setup to breed had high incidences of penile prolapse and clogging of their urogenital tracks with seminal coagulum (five DKOs compared to zero WT male mice), regardless of the female genotype **(Figure 2C)**. These findings, taken together with our protein expression data, prompted a closer investigation into male fitness as the major defect behind the DKO reproductive failure. Indeed, sperm from DKO mice had significantly lower *in vitro* fertilization (IVF) rates compared to wildtype **(Figure 2D)**. Interestingly this may be attributed to sperm fitness / quality rather than quantity as DKO sperm showed trends of decreased motility with no differences in total sperm production **(Figures S2A, S2B and** **2E**).

**Figure 2.**
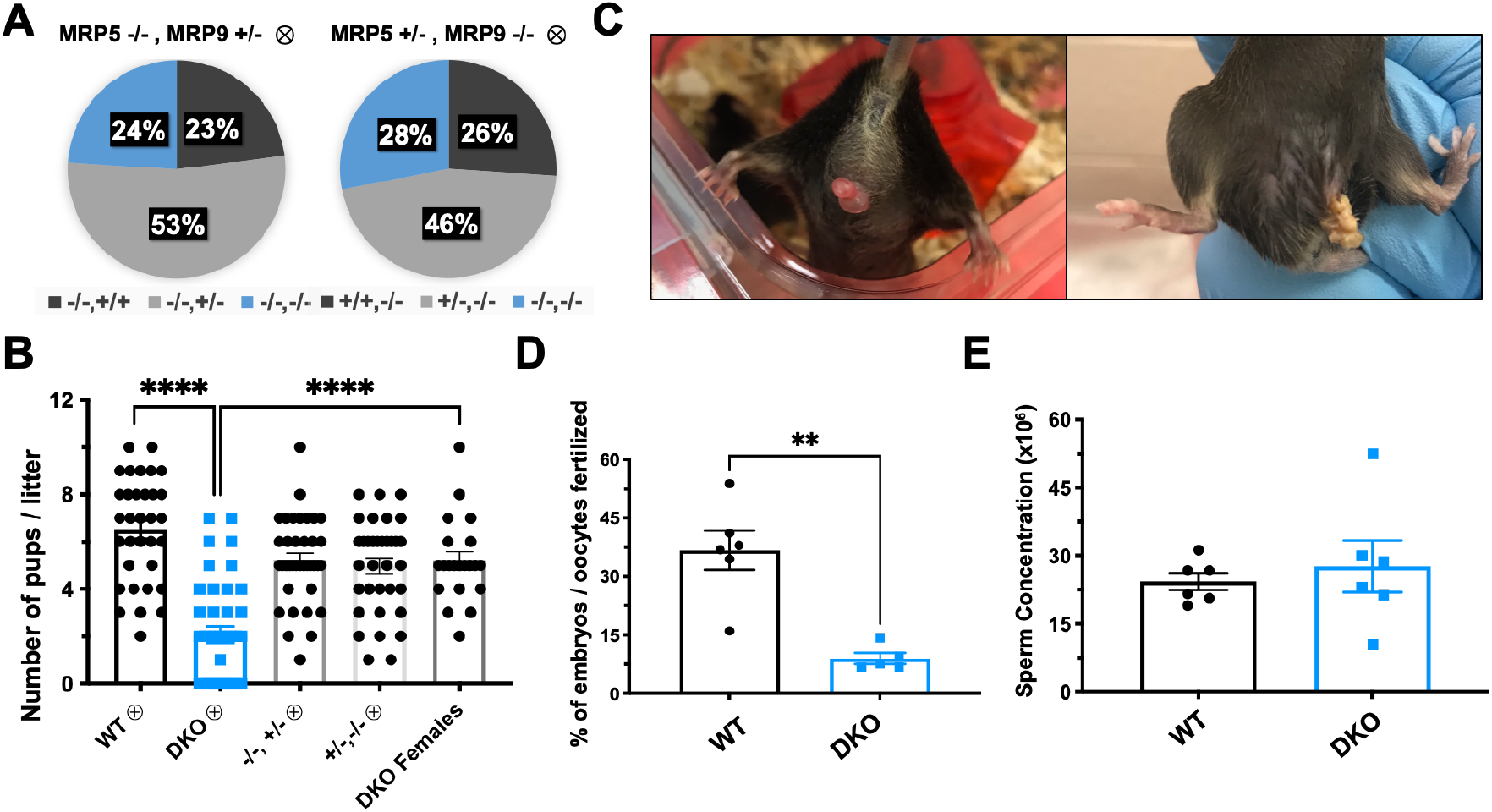
DKO mice are viable but have reduced male reproductive fecundity. **{A}** Genotyped progeny from MRP5−/− and MRR9+/− intercrosses (left), n = 190, Chi-squared = 0.568 and *P* value = 0.753; Genotyped progeny from MRP5+/− and MRR9−/− intercrosses (right), n = 166, Chi-squared = 0.976 and *P* value = 0.614. **{B}** Quantification of the number of pups weaned per litter of intercrossed WT, DKO, MRP5−/− MRR9+/−, MRP5+/− MRR9−/−, and DKO females crossed with WT or double het mice, **** *P* value < 0.0001. **{C}** Representative images of penile prolapse (left) and expressed seminal coagulum (right) in DKO male mice setup for breeding. **{D}** *In vitro* fertilization rate of WT and DKO sperm incubated with WT oocytes for 5 hours at a concentration of 1.0 × 10^6^ sperm/ml. Next day total number of two cell embryos were determined and fertilization rate was calculated based on dividing by the total number of oocytes inseminated for each male, ** *P* value = 0.0019. **{E}** Total concentration of spermatozoa calculated by IVOS system computer assisted sperm analyzer.

### MRP9 is localized to mitochondrial associated membranes in the testes

To understand why DKO male mice have reduced fecundity and sperm fitness we further characterized the function of these proteins in this context. Given their distinct spatiotemporal gene expressions (**Figure S1E**), we hypothesized that these MRPs may play overlapping subcellular roles during spermatid maturation. The Borst group had shown that MRP9 is localized to the sperm midpiece (24, 27, 28), and when expressed in HEK293 closely resembled an ER localization (24). In polarized MDCKII cells, transfection of *ABCC12* showed a distinct intracellular distribution of MRP9 in stark contrast to MRP5 that localized primarily to the basolateral surface of the plasma membrane **(Figure 3A)**. To determine if this was also the case *in vivo* or merely a consequence of ectopic expression, we performed subcellular fractionation of lysates from mouse testes. Immunoblotting showed that MRP9 protein was actually significantly enriched in 9,000 *× g* fractions which pellet mitochondria **(Figure 3B)**. This result was surprising given previously published findings (24, 29, 30), and that MRP9 lacks any obvious mitochondrial targeting sequences. However, an explanation for this unexpected result could be that mitochondrial-associated membranes (MAMs), the contact sites of the ER which are intimately associated with the mitochondria, are enriched in our crude mitochondrial fractions (31–34).

**Figure 3.**
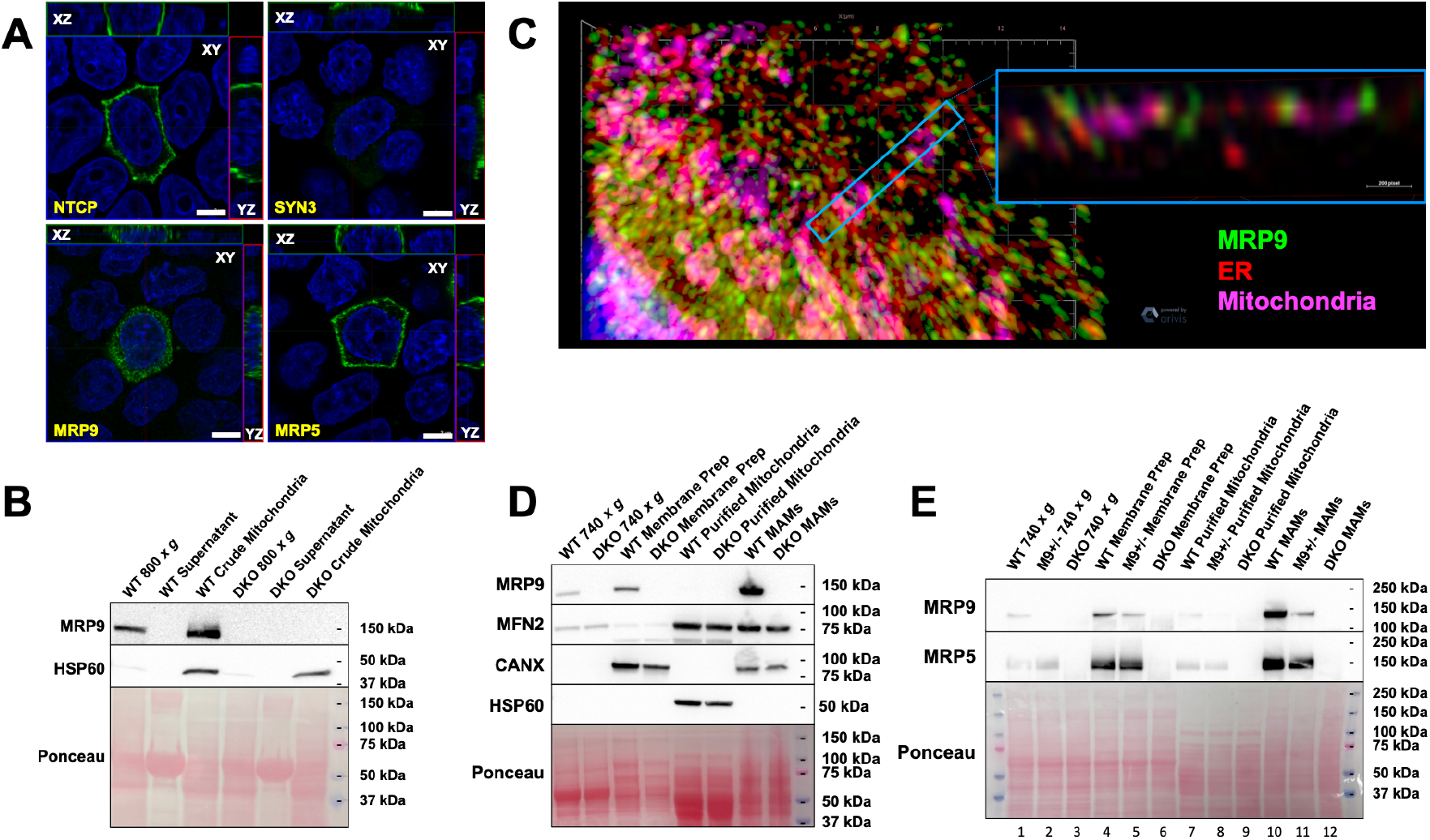
MRP9 is enriched in mitochondrial associated membranes (MAMs) of the testes. **{A}** MDCKII cells grown on transwell filters transfected with basolateral marker sodium taurocholate cotransporting polypeptide *NTCP-GFP*, apical marker syntaxin 3 *SYN3-GFP*, *ABCC12* or *ABCC5*. Polarization of the monolayers was additionally confirmed prior to fixation by measurement of trans-epithelial electrical resistance. A representative confocal section (XY) is depicted along with the composite stacks in the side panel views (XZ and YZ), scale bar equals 5 μm. **{B}** Immunoblot analysis of crude mitochondrial fractionation from testes of mice. Membrane was probed for MRP9 and HSP60, a mitochondrial matrix protein marker. **{C}** Airyscan super resolution microscopy and 3D rendering of HeLa cells transfected with human *ABCC12* for 48 hours and treated with Mitotracker Deep Red FM immediately prior to fixation and immunofluorescence. Antibody probing for MRP9 and Calnexin was followed by Alexa-488 and Alexa-568 secondary antibodies respectively followed by DAPI counter staining prior to mounting. **{D}** Immunoblot analysis of subcellular fractionation from testes of mice. Membranes were probed for MRP9 (M9II-3 antibody) and MFN2, CANX, HSP60 markers of outer mitochondrial membrane, ER membrane, and mitochondrial matrix respectively. **{E}** Immunoblot of subcellular fractionation from testes of wildtype, MRP9 heterozygous and double knockout mice. Membranes were probed for MRP9 (M9I-27 antibody) and MRP5.

Next, we performed Airyscan super resolution microscopy on HeLa cells transiently transfected with human *ABCC12* **(Figure 3C)**. 3D rendering revealed that MRP9 was in close proximity or at the interface of the mitochondria (Mitotracker) and ER (Calnexin) **(Figure 3C and Movie S1)**. To validate this localization, we performed Percoll-based differential centrifugations to further separate crude mitochondrial fractions into purified mitochondria and MAMs from testes tissue homogenates (32). Immunoblots of distinct testes fractions revealed that MRP9 was in fact not present in purified mitochondria but significantly enriched in MAMs, characterized by the markers Mitofusin 2 and Calnexin **(Figure 3D)**. Interestingly, probing for MRP5 from these same fractions indicated that it too was enriched in the MAMs as well as total membranes **(Figure 3E)**. These results indicate that subcellular expression patterns of MRPs may vary dependent on tissue contexts.

### Multi-omics of DKO testes reveals extensive perturbation and mitochondrial dysfunction

In order to systematically dissect why the combined loss of MRP9 and MRP5 manifests in male reproductive dysfunction, we sought to understand what substrates may be transported by these proteins and how their absence may impact homeostatic pathways in the testes. To achieve this, we performed untargeted metabolomics and RNAseq from paired testes of the same mice to integrate any differential metabolites and gene expression changes **(Figure S3A)**. Global untargeted metabolomics of testes indicated significant perturbation, with 24 of the top 25 differential species measured from negative ion mode acquisition of aqueous phase metabolites showing accumulation in the DKO **(Figure 4A)**. Dramatic differences in metabolite profiles were evident however regardless of the extraction method (aqueous vs organic) or the ion mode of acquisition (negative vs positive) **(Figures S3B and S3C)**. Analysis of these data allowed for the identification of mass-to-charge ratios (m/z) of hyper-accumulated candidate metabolites. However, given our sampling was untargeted, a particular *m/z* could represent multiple possibilities of actual biological compounds even given the high mass accuracy and resolution of the Q-TOF platform. As a means to associate these putative species with specific metabolites, all significant m/z’s identified from our six untargeted datasets were pooled and processed using MetaboAnalyst Mummichog analyses (35). From the species of interest analyzed, 65 metabolites were assigned possible KEGG IDs **(Table S1)**. Amongst these, succinyl-CoA, which is also an essential TCA cycle energy source and a heme precursor, had the largest fold-change increase in DKO testes (36, 37).

**Figure 4.**
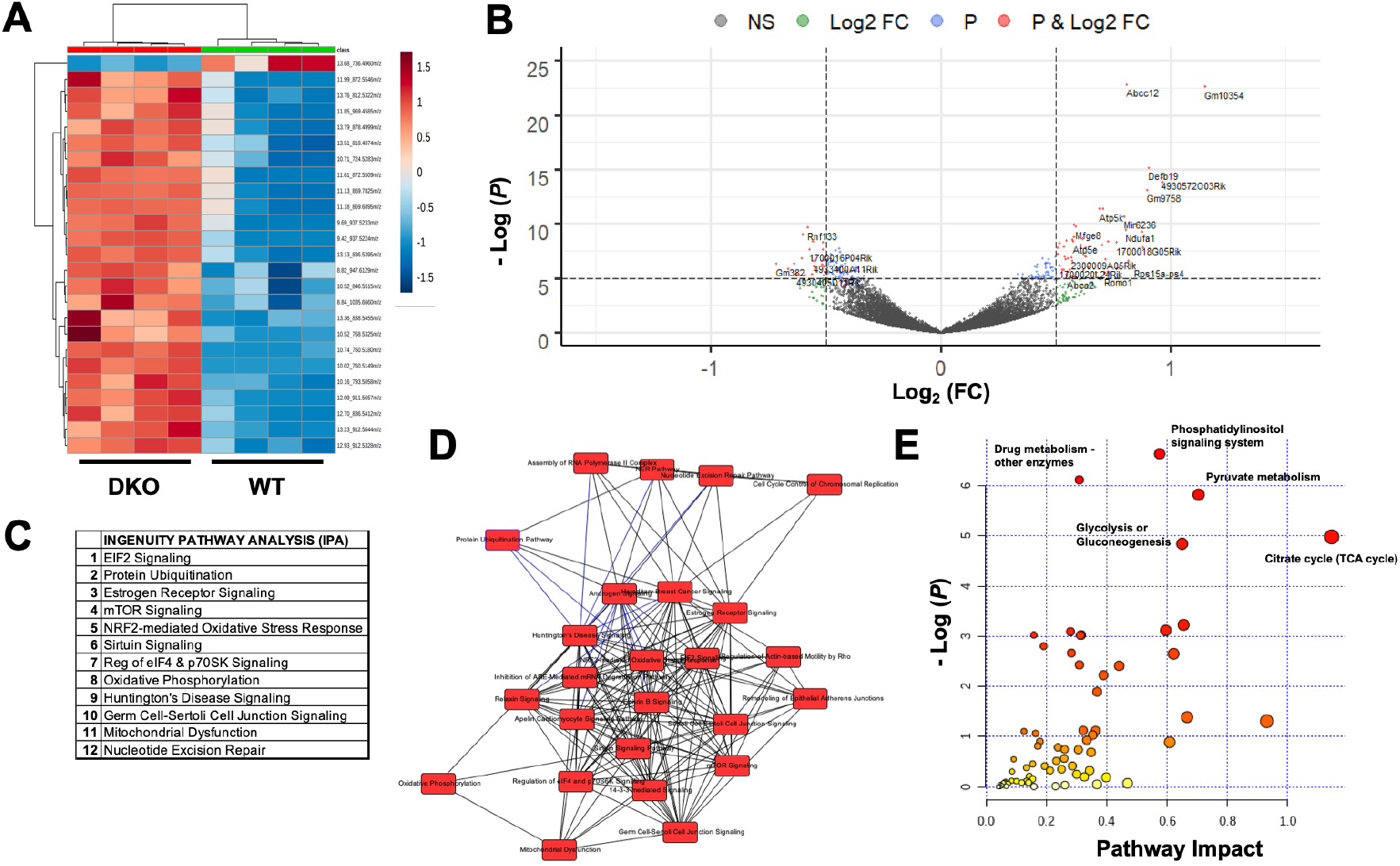
Metabolomics and transcriptomics reveal pervasive perturbation in DKO testes. **{A}** Variable Importance in Projection (VIP) Heatmap of the top 25 differential metabolites in testes from aqueous phase extractions on C18 column run in negative mode**. {B}** Volcano plot of gene expression changes from DKO and WT testes; significance threshold set at −log(*P* value) > 5. **{C}** Top twelve Gene Ontology (GO) Pathways perturbed in DKO testes based on Qiagen IPA analysis. **{D}** Key overlapping canonical pathways in testes based on Qiagen IPA analysis. **{E}** MetaboAnalyst pathway impact analysis from combining RNAseq differential gene expression changes and metabolomics mummichog putative KEGG IDs of the testes; top five pathways by *P* value are annotated.

RNAseq analysis from the same tissues found 3,736 differentially expressed genes between WT and DKO, with a number of top genes differentially expressed including uncharacterized and testis-specific predicted genes (*Gm10354*, *Gm9758*, *5430401F13Rik*) as well as mitochondrial respiratory genes (*Atp5k*, *Ndufa1*, *Atp5e*) **(Figure 4B)**. To unbiasedly determine the pathways and Gene Ontology (GO) terms most significantly affected in these mice we performed Ingenuity Pathway Analysis (IPA) **(Figures 4C and S3D)**. The majority of the top GO canonical pathways identified by IPA were intimately interconnected and overlapped with mitochondrial function **(Figures 4C and 4D)**. Of note, the top two pathways identified, EIF2 Signaling and Protein Ubiquitination, are critical for mitigating unfolded protein response and mitochondrial dysfunction associated with iron and heme deficiencies (38).

To generate metabolite-gene relationships, we integrated our metabolomics and RNAseq data using MetaboAnalyst Joint Pathway Analysis. The merged data analysis revealed significant mitochondrial pathway dysfunction with energy production components of the cell (TCA Cycle, Pyruvate Metabolism, and Glycolysis) showing the highest impact **(Figure 4E)**. Taken together, these findings reinforce a role for MRP9 and MRP5 in mitochondrial metabolism in the testes.

### EIF2a phosphorylation and associated signaling pathways mitigate mitochondrial dysfunction

Having identified mitochondrial dysfunction as the overarching factor from our integrative analysis, we sought to determine the specific mechanisms that underlie this dysfunction in the testes. Mining our RNAseq dataset, we built an analysis pipeline to determine and curate fold changes of all genes associated with a given pathway from published GO databases for downstream investigation and mapping. In querying all genes associated with the mitochondria and mitochondrial dysfunction, we identified over 200 dysregulated genes (**Figure 5A**). We then asked what individual genes or pathways may be involved in responding to such a significant level of disruption. Given the top-related pathway identified from our IPA studies was EIF2 signaling, we investigated all associated genes and found 95 that were perturbed in DKO testes (**Figure 5B**). EIF2 signaling and EIF2a phosphorylation by the heme responsive kinase HRI (*Eif2ak1*) are established mechanisms for regulating translation to reduce mitochondrial stress and unfolded protein response (UPR) in situations of intracellular heme deficiency (38–41). In particular this model is well documented in the erythron, where heme is critical as it becomes the limiting factor for incorporation into hemoglobin for red blood cell differentiation **(Figure S4A**). Limited studies have characterized a similar role in mitigating mitochondrial dysfunction in neurons, but very little is known about HRI and EIF2a signaling in other tissues such as the testes (39, 42–45). Indeed, analysis of the testes single cell RNAseq dataset determined that both factors are expressed during spermatid maturation **(Figure S4B)**, and immunoblotting of total testes homogenates confirmed significant EIF2a protein expression **(Figure 5C)** (26). Importantly, EIF2a phosphorylation was significantly increased in the DKO mice, substantiating the involvement of the EIF2a pathway in mitigating UPR and mitochondrial dysfunction in the testes **(Figure 5C and 5D)**. We next asked if the observed EIF2 signaling in the testes of DKO mice was in response to heme deficiency, as is the hallmark in the erythron. If so, one would expect that iron/heme homeostasis should also be perturbed in the testes of these mice. Subsequent probing with our pipeline pathway revealed heme and iron homeostasis related genes were in fact significantly altered **(Figure S4C)**. To determine if heme levels themselves were also impacted, we measured total heme in male reproductive tissues. Heme levels in the caudal epididymides, the storage sites for mature sperm, showed a significant decrease **(Figure 5E)**, while whole testes and seminal vesicles showed no differences **(Figure S4D and S4E)**. These results suggest that there is insufficient heme specifically in spermatozoa of DKO mice.

**Figure 5.**
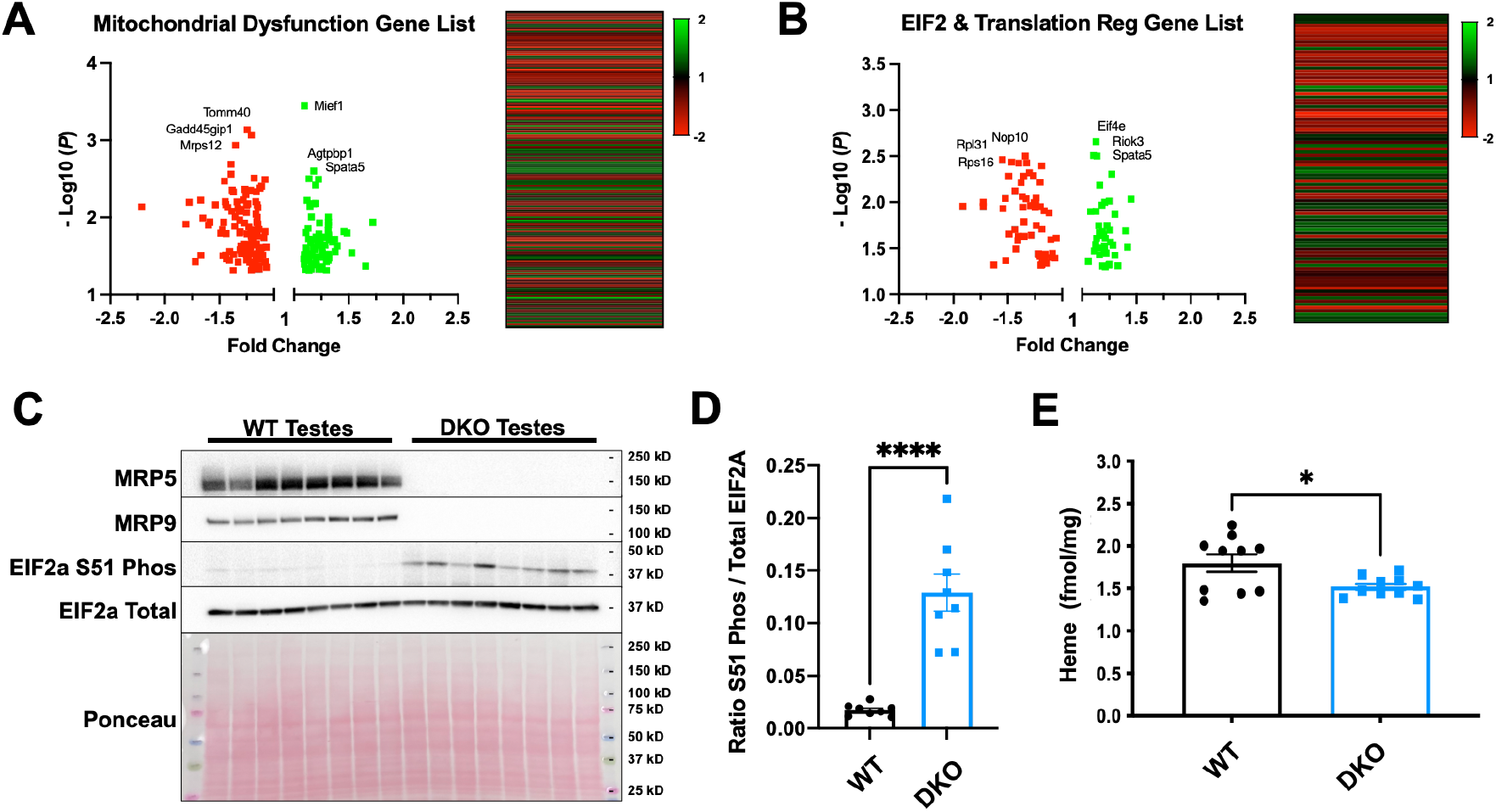
Loss of MRP5 and MRP9 induces EIF2 signaling to mitigate mitochondrial dysfunction. **{A}** Output of pathway pipeline analysis investigating all “mitochondria” and “mitochondrial disfunction” GO pathway gene lists. Gene lists were curated for integration with RNA expression data. Real expression was validated by thresholding a minimum 10 transcripts per million and statistically significant genes (*P* < 0.05) fold change were output in volcano plot and heatmap with 200 total genes identified, sorted by ascending *P* value. **{B}** Output of pathway pipeline analysis investigating all “EIF2 signaling” and “translational regulation” GO pathway gene lists. Gene lists were curated for integration with RNA expression data. Real expression was validated by thresholding a minimum 10 transcripts per million and statistically significant genes (*P* < 0.05) fold change were output in volcano plot and heatmap with 95 total genes identified, sorted by ascending *P* value. **{C}** Immunoblot analysis of testes lysates from DKO and WT mice probed for MRP5, MRP9, Serine 51 phosphorylated EIF2a and total EIF2a protein. Blot is representative of three separate experiments, each lane represents an individual mouse, n=8 mice per genotype. **{D}** Quantification of DKO and WT phosphorylated EIF2a normalized to total EIF2a protein in subpanel C, **** *P* value < 0.0001. **{E}** Heme content of pooled caudal epididymides from DKO and WT mice by oxalic acid quantification, n=15 animals per genotype (1.5 mice or 3 epididymides per replicate), * *P* value = 0.0275.

**Figure 6.**
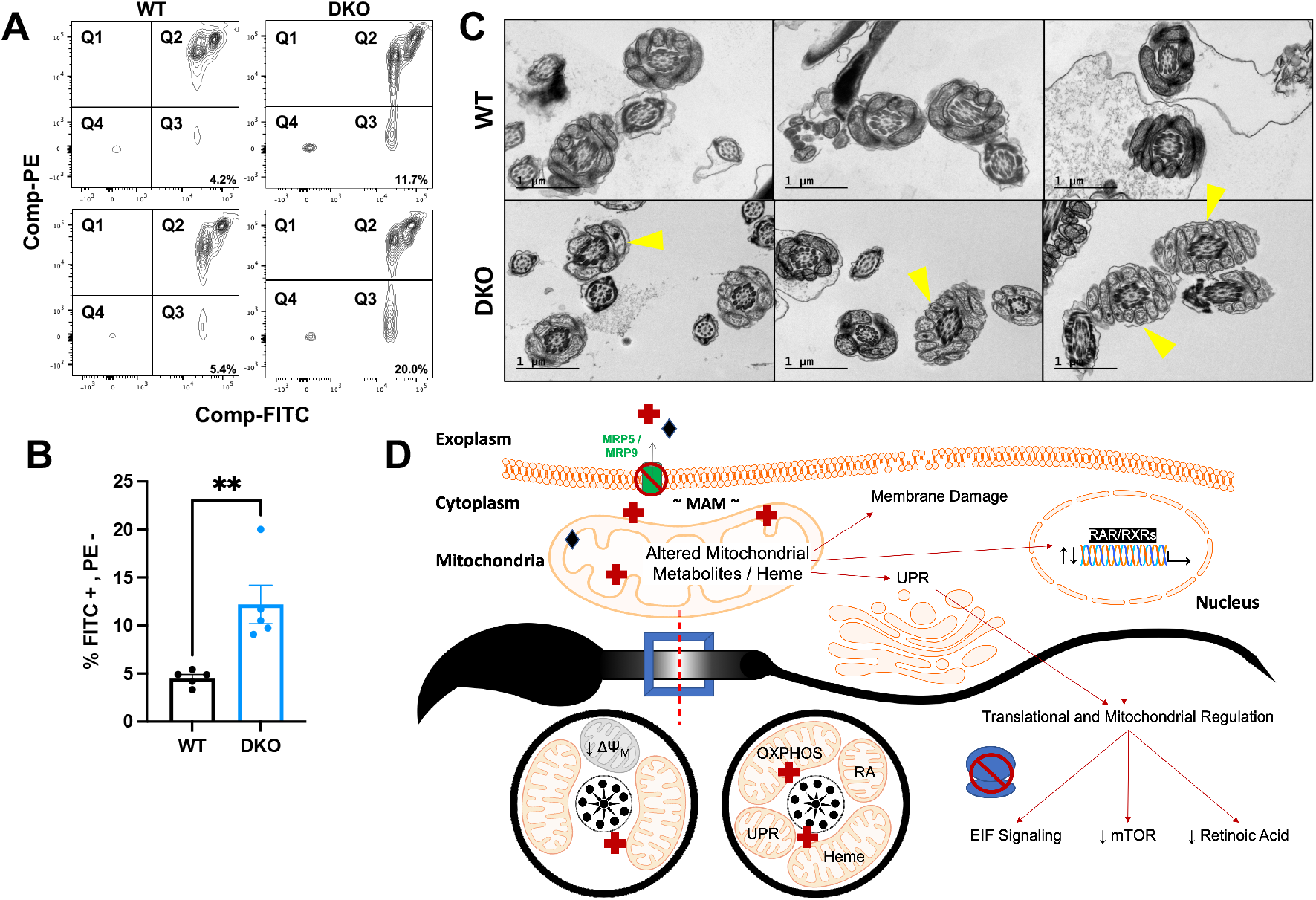
Spermatozoa are severely compromised by the combined loss of MRP5 and MRP9. **{A}** Representative sorting analysis of WT and DKO sperm stained with JC-1. Q3 is gating for populations that are FITC-pos and PE-neg representing a reduction J-aggregates / red fluorescence and therefore decreased mitochondrial membrane potential. **{B}** Quantification of sperm stained with JC-1 and gated for populations that are FITC-pos and PE-neg (from subpanel 6a), n=5 animals, ** *P* value = 0.0055. **{C}** TEM of swum-out caudal epididymal spermatozoa from WT (top) and DKO (bottom). Cross sections of the sperm midpiece visualize cristae and mitochondrial morphology of the mitochondrial sheath. Yellow arrow heads highlight vacuolated and aberrant mitochondria. Representative images from at least 15 FOVs per sample, scalebar equal 1 μm. **{D}** Model figure – the combined ablation of MRP9 and MRP5 causes inappropriate trafficking or distribution of metabolites such as heme and/or other substrates inducing mitochondrial damage. This dysfunction induces signaling through the EIF2 and mTOR pathways to regulate protein translation and UPR with corresponding changes in retinoic acid levels and signaling. Germ cells differentiate into mature spermatozoa, but these are functionally defective as they fail to fertilize normal oocytes due to mitochondrial insufficiencies.

A critical component of the canonical EIF2 signaling cascade for maintaining mitochondrial function and redox homeostasis is the alteration of mTOR signaling to inhibit proliferation and enable differentiation (38, 46). Indeed, the mTOR signaling pathway was one of the most significant GO terms identified by IPA **(Figure 4C and S3D)**, which we also validated with our analysis pipeline **(Figure S5A)**. Upregulation of EIF-4E, a highly abundant translation regulation factor in the testes, has been shown to play an important role in spermatogenesis through regulation of capping stage-specific mRNAs during germ cell development (47). A modulator of these related pathways is retinoic acid, which has been implicated in mTOR/EIF-4E cascade and plays a key role in a variety of biological contexts including regulation of genes impacting mitochondrial function (48–52). Principal targets of retinoic acid include key OxPhos and respiratory genes such as NADH dehydrogenase subunits, cytochrome c oxidases and the ATPases, some of the top differentially expressed genes from our RNAseq **(Figure 4B)** (51, 53). Investigation of retinoic acid related genes revealed significant perturbation including decreases in the cytosolic chaperone *Crabp1*, responsible for delivering retinoic acid for catabolism, as well as binding receptors (RAR, RXRs), suggestive of dysregulated retinoic acid homeostasis **(Figure S5B)** (54–56). Importantly, retinoic acid signaling is essential for male germ cell differentiation, exerting key control over meiotic initiation and spermatogenesis, with defects in retinoid homeostasis a well-established cause of male infertility (57–60). Taken together, we hypothesized that retinoic acid homeostasis/signaling may play an important role in the phenotypes observed in our DKO mice. Quantification of the major components of the retinoid pathway revealed that total levels of vitamin A (retinol) and storage esters were not perturbed in the testes **(Figure S5C and S5D)**, but all-*trans* retinoic acid levels were significantly reduced DKOs compared to wildtype **(Figure S5E)**. Other retinoic acid isomers were unchanged (13-*cis* retinoic acid, 9,13-di-*cis* retinoic acid) or below the limit of detection (9-*cis* retinoic acid) (data not shown).

To further validate if retinoic acid binding transcription factors and cascade regulators were indeed impacting mitochondrial genes of interest, we performed unbiased motif discovery on regulatory sequences of all up-regulated and down-regulated genes from our pathway pipeline **(Figure 5A and S5F)**. The top five statistically significant DNA sequences identified were further analyzed and queried against known binding motifs. With two independent analysis platforms (MEME suite and HOMER) we identified significant enrichment of conserved retinoic acid related binding motifs including putative RXRs and RORA binding sequences **(Figure S5F)** (61). These findings suggest a mechanistic link between transcripts related to mitochondrial dysfunction, retinoid signaling genes, and decreased retinoic acid levels, corresponding with a specific enrichment of retinoic acid binding motifs in the DKO mice.

### Ablation of MRP9 and MRP5 significantly damages sperm mitochondria

To determine whether mitochondrial disruption could be inducing the gene expression and metabolomic changes that in-turn impact sperm fertilization in the DKO testes, we analyzed mitochondria oxidative phosphorylation complexes in total testes lysates. However, all five complexes appeared to be normal and unchanged in DKOs **(Figure S6A)**. We next fractionated the testes and performed high-throughput targeted metabolomics on each isolated fraction, allowing for the identification and quantification of specific metabolites within each subcellular niche **(Figure S6B)**. We found that the mitochondria and MAM fractions of the DKO testes hyper-accumulate triglycerides, indicative of mitochondrial metabolism dysfunction **(Figure S6C and S6D)** (62, 63).

Mature caudal epididymal sperm were then collected from WT and DKO adult mice by swim-out for mitochondrial membrane potential (ΔΨ_M_) analysis and transmission electron microscopy (TEM). Staining of sperm with JC-1 revealed that DKO mice have reduced J aggregates and ΔΨ_M_ compared to WT **(Figure 7A, 7B and S7A)**. Strikingly, TEM cross sections of sperm midpieces revealed DKO mice also have highly vacuolated and aberrant mitochondria, in contrast to the ubiquitously electron dense cristae in the WT **(Figure 7C)**. These defects were not observed in the single KOs or double heterozygous animals **(Figure S7B)**.

## Discussion

ABCC proteins are one of the most well-studied class of transporters due to their critical roles in cancer biology, multidrug resistance and associated human diseases (3, 64–67). Taken together, our studies fill significant gaps in our understanding of not only of MRP9 but also MRP5, uncovering novel endogenous functions related to reproduction. Given MRP5 is genetically dispensable in the mouse model with low gene expression in male reproductive tissues, no studies have examined its protein levels or function in the testes (14, 68–70). Klein et al showed a weak MRP5 signal in Leydig cells of rats but nothing in macaque or human testis (71). We show that MRP5 is detectable in the testes, but that it also appears to be enriched in mitochondrial associated membranes along with MRP9. Though MRP5 still localizes to basolateral plasma membranes in polarized MDCKII cells, our tissue fractionation studies bring to light the possibility of additional novel tissue dependent expression patterns and subcellular localizations for this well studied transporter. We envision that these results may have significant implications when it comes to cancer and the distinct roles MRPs may play dependent upon tissue / cell type. We propose the following model for the concerted function of MRP9 and MRP5 in the testes **(Figure 7D)**: In the absence of MRP9 and MRP5, metabolites such as heme and/or other substrates, are inappropriately trafficked or distributed causing mitochondrial damage. This dysfunction induces signaling through the EIF2 and mTOR pathways to regulate protein translation and UPR with corresponding changes in retinoic acid levels and signaling. Germ cells differentiate into mature spermatozoa, but these are functionally defective as they fail to fertilize normal oocytes due to mitochondrial insufficiencies.

MRP9 is an enigmatic ABCC transporter. It has a unique expression pattern, no known drug substrates, and the last in the ABCC class still to be characterized (6, 24, 25). In the current study, we fill in some of the gaps in our knowledge by generating an MRP9 knockout mouse model to explore its endogenous functions. We find that MRP9 is specifically expressed in developing spermatids and mature spermatozoa in mice and unlike most other MRPs, MRP9 does not localize to the plasma membrane irrespective of cell polarization, consistent with the published findings of Ono et al (24). Contrary to other reported studies however, MRP9 is not endogenously expressed in any cell lines tested in our hands, including primary mouse embryonic fibroblasts, and MRP9 protein cannot be induced by sodium butyrate treatment **(Figure S8)** as was reported recently (72).

Transient transfections and tissue fractionations provide strong evidence that MRP9 partitions specifically to mitochondrial-associated membranes of the testes, a distinct compartment characterized by mitochondrial outer membrane and endoplasmic reticulum markers. This is physiologically reasonable as MRP9 is localized to developing and mature spermatozoa which are known to be devoid of traditional ER structures (73, 74). Though the ER is depleted in the process of differentiation, vestiges of ER membranes likely remain, particularly those in tight contact with mitochondria, and have been postulated to concentrate in the neck of the sperm midpiece to serve as a store for Ca^2+^ necessary for sperm activation (74, 75). Indeed, ER resident proteins such as ERp29 and Calreticulin have also been demonstrated to be present in sperm and play essential roles in acrosome reaction, fertilization and stress responses (74, 75). We envision MRP9 may reside at these interfaces adjacent to mitochondria of the midpiece, effluxing substrates essential for regulating mitochondrial function and/or trafficking metabolites between these compartments as spermatogonia develop.

Loss of MRP9 alone has no discernable impact on male reproductive function and knockout mice are viable. Additionally, histological inspection of the testis and epididymis of these mice reveal effective spermatogenesis, with normal spermatid progression in seminiferous tubules and mature sperm in their convoluted ducts. Unsurprisingly, we also see no morphological defects evident by electron microscopy of sperm from MRP9 single knockouts, as these male mice are perfectly capable of siring progeny even in a MRP5 heterozygous background. Any discernable impact on reproduction is only found when both MRP9 and MRP5 are ablated. Indeed, DKO mice show significant reproductive failures due to decreased male fecundity. Defects in sperm fertilization rates and high incidences of penile prolapse and seminal coagulum clogging contribute to their decrease in reproductive output as measured by litter size. This is likely due to the DKO sperm’s inability to effectively fertilize, as supported by the drastically reduced IVF rates.

In addition to decreased motility trends, we show substantial changes in the composition of metabolites retained in the testes as well as significant perturbations in gene expression as a consequence of the loss of MRP9 and MRP5. Importantly, our integrative analysis of these approaches demonstrate that energy production and mitochondrial function were the most impacted pathways in the DKO testes, consistent with our localization studies. Targeted high-throughput metabolomics of subcellular fractions also revealed substantial changes at the subcellular level. While others have shown that triglyceride species can accumulate at the whole tissue level during mitochondrial dysfunction, we show that these lipid changes are occurring at the mitochondria and MAMs in the DKO testes (62, 63, 76, 77).

The top pathway identified by our gene expression analysis was EIF2 signaling, which we validated *in vivo* as a response to mitochondria dysfunction and/or induced by alterations in heme levels in maturing sperm. Indeed, heme and iron related genes were differentially expressed in the testes, and quantification of heme in the caudal epididymis of DKO mice showed significantly decreased levels. In a similar vein to the well-established models of unfolded protein response in the erythron, we envisage a necessity for heme sharing and partitioning as mitochondria are rapidly being recruited and cells are undergoing proliferation for spermatogenesis and differentiation. Though this translational regulation via EIF2 signaling has been well-studied in other contexts, virtually nothing is known about its role in the testes. The only published study to our knowledge relates to EIF2s3y, which was shown to be essential for regulating protein proliferation in spermatogenesis (78). It is yet to be determined whether or not this subunit plays any direct role to our findings here, although it is also significantly upregulated in the DKO testes (*P* = 0.022).

Beyond EIF2 signaling, IPA identified several additional pathways: Protein Ubiquitination (essential for unfolded protein response), mTOR Signaling, Regulation of EIF4, Oxidative Phosphorylation, Germ Cell-Sertoli Cell Junction Signaling (an essential component of sperm differentiation), and Mitochondrial Dysfunction. It is plausible that all-*trans* retinoic acid may be a regulator in all these pathways for the following reasons: 1) it is essential in male fertility as one of the primary signaling molecules from Sertoli cells for progression of spermatogenesis; 2) it is critical for regulating gene expression of essential oxidative phosphorylation and mitochondrial genes; and 3) it has been shown to modulate mTOR and EIF4 regulation (51, 52, 57). Consistent with this notion, we find genes associated with retinoids and retinoic acid levels significantly decreased in DKO testes. Furthermore, targets of gene expression perturbation have significant enrichment of retinoic acid-related binding motifs for transcriptional regulation, albeit modulation via these motifs may be indirect. RORA has been reported to act as a constitutive activator of transcription in the absence of ligand and all-*trans* retinoic acid does not bind to RXR with high affinity as compared to 9-*cis* retinoic acid (79, 80).

These systemic changes in the testes of DKO mice are in-line with the striking morphological abnormality of their mitochondria at the ultrastructural level. The impact of such evident dysfunction at the sperm midpiece has significant effects on male fecundity. The mitochondrial sheath is essential for reproductive fitness as these tightly wrapped mitochondria are critical for ATP production and maximize efficiency of energy delivery, directly correlated with sperm swimming velocities across species (27, 28, 81–84). However, this immense output comes with a price - because spermatozoa are so densely packed with mitochondria, they are also epicenters for the production of detrimental free radicals/reactive oxygen species (27, 81, 83, 85). If not balanced and mitigated, this can cause myriads of problems including lipid peroxidation, which has been shown to damage the sperm and decrease their mitochondrial membrane potential (27, 81, 83, 86). Loss of MRP5 and MRP9 alters mitochondrial homeostasis not only inducing vacuolated and aberrant mitochondria, but also resulting in a significant reduction in sperm mitochondrial membrane potential as observed by JC-1 staining. This relationship between mitochondrial membrane potential and sperm function directly correlates with mitochondrial health, sperm motility, and modulation of respiratory chain activity and oxidative phosphorylation pathways (27, 82, 87–89). Concordantly, male fecundity relies heavily on this organelle and its membranes being fully intact and functional for the overall health of the spermatozoa and their fertilization capacity, cementing why DKO males show such significant loss of biological function (27, 81–83). Male infertility is a significant problem globally, affecting over 70 million people and accounting for 50% of documented issues in infertile couples (82, 90, 91). To our knowledge, mutations or SNP variants of MRPs have not yet been investigated for any correlation to cases of male reproductive dysfunction. Our findings here would suggest that human patients with infertility issues, typically assayed for and characterized by mitochondrial dysfunctions, may have hypomorphic mutations in MRP9 and MRP5.

## Materials and Methods

### General methods

#### Animals

All mice used were housed in Animal Sciences Departmental facilities at the University of Maryland and kept in standard 12-hour light-dark cycles. Both sexes were used for all experiments unless stipulated otherwise, such as the analysis of male sex organs and requisite male reproduction phenotypes. ABCC12/MRP9 mutant mice were generated via CRISPR/Cas9 RNA injections at the National Human Genome Research Institute (NHGRI). Guide RNAs **(Table S3)** were purchased from Sage Laboratories, St Louis, MO and Cas9 RNA was purchased from Trilink Biotechnologies, San Diego, CA. The guide RNA and Cas9 RNA were combined at a concentration of 5 ng/μl each in 10 mM Tris, 0.25 mM EDTA pH 7.5 for pronuclear injection. Performed using standard procedures, fertilized eggs were collected from super-ovulated C57BL/6J females approximately nine hours after mating with C57BL/6J male mice. Pronuclei were injected with a capillary needle with a 1-2 μm opening pulled with a Sutter P-1000 micropipette puller. The RNAs were injected using a FemtoJet 4i (Eppendorf) with continuous flow estimated to deposit approximately 2 pl of solution. Injected eggs were surgically transferred to pseudo-pregnant CB6 F1 recipient females. DNA was then obtained from founder (F0) animals by tail biopsy and sequenced to determine mutations. F0 animals carrying mutations were crossed to C57BL/6 animals and the resulting heterozygous F1 animals were either intercrossed to generate homozygous mutant animals or back crossed to C57BL/6 mice for propagation. MRP5 mutant mice, a gift from Dr. Piet Borst (Netherlands Cancer Institute), had previously been outcrossed into full C57BL/6 background and were bred to MRP9 mutant mice to generate double knockouts (DKOs). MRP5 and MRP9 mice were genotyped from tail genomic DNA via PCR and RFLP methods (see **Table S4** for complete primer sequences). In brief, tail snips were taken from weaned 21-day old (P21) pups and digested. The MRP5 allele was determined as described previously via amplification of specific MRP5 and hygromycin cassette primers (14). MRP9 mutants having a +17bp insertion were PCR amplified and subsequently restriction enzyme digested with AvaII (specific to the mutation). All subsequent protocols described below involving these animals were approved by the Institutional Animal Care and Use Committee (IACUC) at the University of Maryland, College Park.

#### Standard Immunoblotting Methods

For specific detection of MRP5 and MRP9 protein lysates, as well all other general proteins unless otherwise specified, the following protocol was used: Lysis in lysis buffer (0.75% NP40, 0.5% Triton X-100, 0.1% SDS, 1.75 mM EDTA, 1.25 mM EGTA, 0.25% DOC, 150 mM NaCl, 50 mM HEPES, 25 mM Tris-Cl pH 7.5) with 3x protease inhibitors (cOmplete Protease Inhibitor Cocktail EDTA-free - 1 mM phenylmethylsulfonyl fluoride, 4 mM benzamidine, 2 μg/ml leupeptin, and 1 μg/ml pepstatin, Roche Diagnostics, cat. number 1187358001) and 3x phosphatase inhibitors (Millipore Inhibitor Cocktail Set, cat. number 524627) and sonicated twice. Protein homogenates were then clarified via centrifugation at 15,000 *× g* for 15 min at 4°C and insoluble pellets discarded. Total protein concentration of the supernatant was measured using the Bradford assay (Bradford Reagent Protein Assay Dye Reagent, Bio-Rad, cat. number 5000006). Unboiled protein samples were then mixed with 4x Laemmli sample buffer and 100 mM dithiothreitol (DTT), and 100 μg of protein/lane was loaded and separated in an SDS-PAGE gel and transferred to a nitrocellulose membrane. Proteins were then cross-linked to membranes by UV treatment and stained with ponceau S solution for 3 minutes for protein loading control before incubation in blocking buffer (5% nonfat dry milk in 0.05% PBS-Tween 20 [PBS-T]) for 1-hour rocking at room temperature. Blots were then incubated overnight at 4°C in blocking buffer containing primary antibody of interest; anti-MRP5 and anti-MRP9 rat monoclonal antibodies are always used at a concentration of 1:200 unless specified otherwise. The following day blots are rinsed once quickly (2 min) and then washed three times (15 min) in PBS-T and subsequently incubated for one hour with horseradish peroxidase (HRP)-conjugated goat anti-rat IgG secondary antibody at 1:20,000 (Invitrogen cat. number 31470). Blots were then rinsed again once quickly, followed by three washes (20 min) with PBS-T and then developed using enhanced chemiluminescence (SuperSignal West Pico PLUS Reagent, Pierce, cat. number 34580) and detected using ChemiDocTM Imaging Systems (Bio-Rad). If subsequent re-probing was necessary, blots were then stripped with 500 mM NaOH for 10min and rinsed quickly with DI H_2_O twice, and then PBS-T three times (5min) before blocking again for one hour to repeat the process with subsequent antibodies.

#### Subcellular fractionation and MAM isolation

Mitochondrial Associated Membranes (MAMs) isolation and subcellular fractionizations were based off of Wieckowski et al 2009 Nature Methods Protocol with minor modifications for optimization of testes tissue. Age matched wildtype and double knock out male mice were anesthetized with intraperitoneal injection of ketamine/xylene and then systemic perfusion was performed via standard cardiac puncture method. Perfusion was performed with IB-1 buffer (225 mM mannitol, 75 mM sucrose, 0.5% BSA, 0.5 mM EGTA and 30 mM Tris–HCl pH 7.4) using 25 ml total pushed through a 30 ml syringe with a 26G needle (BD Precision Glide, cat. number 305111) into the left ventricle of the beating heart, which upon completion resulted in completely cleared liver and kidneys. Livers and testes were then immediately harvested and placed in IB-3 buffer (225 mM mannitol, 75 mM sucrose and 30 mM Tris–HCl pH 7.4) on ice at 4°C and the remaining isolation was performed entirely at 4°C in a cold room. A single liver or four pooled testes were necessary to achieve sufficient yield of purified MAMs, which were minced and combined and then lysed in IB-1 buffer via 10 strokes in a 22 gauge 5ml glass duall dounce homogenizer. The homogenate was then spun down twice at 740 *× g* to remove nuclei and cellular debris. The remaining supernatant was spun at 9,000 *× g* for the isolation of crude mitochondria, which was subsequently washed and spun down twice at 10,000 *× g*. The first 10,000 *× g* spin was in IB-2 buffer (225 mM mannitol, 75 mM sucrose, 0.16% BSA and 30 mM Tris–HCl pH 7.4) and then the second in IB-3 for further purification prior to a 100,000 *× g* spin through a Percoll (Sigma, cat. number P1644) gradient to separate MAMs from mitochondria. The original supernatant from the 9,000 *× g* spin was also spun down at 100,000 *× g* to concentrate the remaining smaller organelles, plasma membranes and cytosolic proteins labeled “Membrane Prep.” Each of the isolated fractions “740 *× g*”, “Membrane Prep”, “Mitochondria” and “MAMs” as well as total initial homogenate were subsequently lysed in lysis buffer and following SIM, ran side by side in a Criterion Precast Gradient 4-15% SDS-PAGE gel for western blotting probing of key intracellular organellar markers and proteins of interest: HSP60 (Abcam, cat. number ab46798), Calnexin (Abcam, cat. number ab22595), Mitofusin 2 (Invitrogen 7H42L13, cat. number 702768), MRP5 and MRP9.

#### Histology and immunohistochemistry

Paraffin-embedded tissues were sectioned at 4 μm and processed for immunohistochemistry (IHC) as previously described in Pek et al with minor modifications (92). Slides were first rehydrated in descending gradient from pure xylene through ethanol series (100%, 95%, 80%, 70%) and followed by endogenous HRP quenching before heat-induced epitope retrieval in citrate buffer pH 6 (Target Retrieval Solution, DAKO, cat. number S1700). After epitope retrieval, sections were then rinsed and blocked with avidin, biotin and protein blocking following Vector labs ABC elite staining kit (Vector labs, cat. number PK-6104) prior to incubation with monoclonal rat anti-MRP9 antibodies at 1:50 overnight at 4°C. The next day sections were then incubated with secondary biotinylated anti-rat antibody for 30 minutes at room temperature. ABC tertiary reagent was added and then signals were detected by DAB substrate incubation per manufacturer recommendations (DAB/Metal Concentrate Peroxidase Kit. Thermo Fischer, cat. number 1859346 and 1856090). Finally, slides were lightly counterstained with hematoxylin and then dehydrated in opposite ascending series and mounted with DPX Mountant (Sigma, cat. number 06522-100).

#### Heme quantification

To extract heme from perfused mouse tissues, a minimum of 30 mg of caudal epididymides (either three or four pooled together), an entire testis or seminal vesicle were dounce homogenized fresh in a minimum of five volumes of lysis solution containing 50 mM Tris/HCl pH 8.0, 5 mM CaCl_2_, 50 mM NaCl, 1% Triton X-100 and 0.5% Proteinase K. Proteinase K was added fresh to the buffer and the homogenate was incubated overnight at 37°C with moderate shaking. The next day the Proteinase K digest is sonicated for 1 min (10 W, pulse 0.5 sec), centrifuged at 11,000 *× g* for 45 minutes and then the supernatant was collected for heme quantification. 20 μl of the tissue lysate sample is mixed with 180 μl of 2 M oxalic acid and sample tubes are then heated in a water bath set at 100°C for 30 minutes while a duplicate set of samples remain at room temperature as blank controls. Fluorescence of porphyrins were then read using Ex400/10nm, Em645/40nm on a BioTek microplate reader (BioTek, Synergy HT) and heme content calculated based off of a heme standard curve and subtraction of blank values (parallel unheated samples in oxalic acid) from samples.

#### EIF2α phosphorylation quantification

Eight age matched wildtype and double knock out male mice were sacrificed by cervical dislocation, and whole testes were dissected out and rinsed in ice cold PBS. A single testis was then homogenized in 500 μl standard lysis buffer with 5x protease inhibitors and 3x phosphatase inhibitors, via 8 strokes in a 20 gauge 2 ml glass duall dounce homogenizer, transferred to 1.5 ml Eppendorf tube and placed on ice for 15 minutes, vortexing for 15 seconds every 5 minutes. Afterward, the homogenate is then sonicated for 10 seconds, twice, prior to proceeding with standard clarifying spin for standard immunoblotting. For all samples, 150 μg of protein was then separated on Criterion Precast Gradient 4-15% SDS-PAGE gels (Bio-Rad, cat. number 5671083) and probed for phosphorylated EIF2α protein levels with rabbit anti-Eif2a S51 phosphorylation polyclonal antibody (1:500 dilution) (Cell Signaling Technologies, cat. number D9G8) and Total EIF2α protein levels with rabbit anti-EIF2a polyclonal antibody (1:10,000 dilution) (Cell Signaling Technologies, cat. number D7D3). Additional blots were probed with rat anti-MRP5 NC3 monoclonal antibody (1:200) and rat anti-MRP9 M9I-27 monoclonal antibody (1:200). Quantification of EIF2α S51 phosphorylation was performed by Image Lab 6.1 software (BioRad) and normalized to total EIF2α protein.

### Mouse reproduction studies

#### Breeding schema for general fecundity and fertility

All breeding studies were performed in harem setups with one stud male mouse and two female mice per cage. These harems were monitored over a period of at least 6 months to track breeding of ABCC5/ABCC12 double knockout mice (−/−, −/−) as well as the requisite control intercrosses of heterozygous-homozygous (+/−, −/−) and homozygous-heterozygous (−/−, +/−) mice. Once successful pregnancies were determined, females were separated from males and individually housed for monitoring fecundity. New female mice were then rotated in the male mice cages to maintain setups of two female mice per stud male. Males unable to produce any pregnancies after two months were removed and replaced. Pregnant dams were monitored at least twice daily for birth of pups and overall health, litter size was recorded at birth and any subsequent mortality logged through weaning was reported. Fecundity was calculated based off the number of breeding setups required to achieve same number of pregnancies as well as the number of offspring per litter and average productivity of litters per harem breeding setup. All genetic segregation and statistics for MRP5 and MRP9 alleles were calculated on weaned P21 mice.

#### *In vitro* fertilization (IVF) and general sperm characterization

All IVF was conducted in HTF media (EmbryoMax, Sigma, cat. number MR-070), supplemented with 3 mg/ml of BSA. Males were mated with females and assessment of mating was determined by the presence of a vaginal plug. Males that were listed as plug positive were allowed to rest a total of three days prior to IVF. Spermatozoa was collected from individual males and placed in 90 μl of TYH media supplemented with beta-methyl-cyclodextrin to induce capacitation (93). Sperm samples were allowed to capacitate for a total of 45-60 minutes. A 10 μl sample was diluted in 1400 μl of IVF media and assessed on an IVOS system (Hamilton Thorne CASA - Computer Assisted Sperm Analyzer). Total sperm concentration, percent progressive motility and percent rapid motile cells was calculated. Visual assessment of morphology was also assessed, and IVF was conducted and standardized to an insemination dose of 1.0 × 10^6^ sperm/ml for each male processed. Oocytes and sperm were incubated for a total of 5 hours. Presumptive zygotes were rinsed through HTF media to remove accessory sperm and cultured overnight in KSOM media plus amino acids (94). The next morning, total number of two cell embryos were determined and a fertilization rate calculated based on the number of two cell embryos per total number of oocytes inseminated.

#### Sperm “swim-out” collection

Age matched male mice were sacrificed by cervical dislocation, and entire reproductive tracts were dissected out intact to isolate the caudal epididymis. Caudal epididymides (CE) were trimmed away from fat pads and the testes, rinsed once quickly (dipping) in DPBS and then dabbed dried (single touch) on a kimwipe before being placed in a 35 mm dish with mineral oil. Prewarmed DPBS with added 100 mg/L MgCl_2_ and CaCl_2_ (swim-out solution) to 37°C was then added to the mineral oil in close proximity to the CEs for sperm swim-out. Swim-out was achieved by snipping the CEs with dissecting micro-scissors and dragging the exposed globule with a 22G needle from the mineral oil into the swim-out solution. The entire petri dish was then covered and then placed in the 37°C incubator for 30min to allow for purified sperm to fully swim-out into the solution. Then sperm were collected with a 1ml pipettor and spun down gently in a 1.5 ml Eppendorf tube at 333 *× g* for 3 minutes, followed by resuspension in fresh swim-out buffer and rocked gently for 2 minutes. This rinsing was repeated the same way again twice, and after the third rinse and spin down the sperm are ready for downstream analysis.

#### Sperm transmission electron microscopy (TEM) imaging

Swim-out collected sperm were resuspended in a final 2% PFA and 1% Glutaraldehyde solution for fixation, and then rinsed in 2 changes of 0.1 M Sodium Cacodylate buffer with 2.4% sucrose and 8 mM CaCl_2_. Samples were then postfixed in 2% Osmium Tetroxide in 0.1 M Sodium Cacodylate buffer with 2.4% sucrose and 8 mM CaCl_2_. Samples were rinsed thoroughly with distilled water, and then en bloc stained in saturated aqueous Uranyl Acetate. Samples were then dehydrated through a graded ethanol series (50%,70%, 95%, 95%, 100%, 100%, 100%, 100%). Samples were transitioned into absolute Acetone for 3 changes. Then infiltrated in a 1:1 (plastic: acetone) and then 3:1 (plastic: acetone) dilutions of Embed 812 and followed with 3 changes of 100% plastic prior to embedding. Samples were cure 24 hours in a 60°C oven. Once removed from the oven samples were thick sectioned with glass knives to determine best area to thin section, on a Leica EM UC6 Ultramicrotome. A diamond knife (Diatome) was used to thin section samples and sections were placed on 150 hex copper grids. Grids were stained sequentially with Aqueous Saturated Uranyl Acetate followed by Reynold’s Lead Citrate. Sections were examined at 120 KV on a JEOL 1400 plus Transmission Electron Microscope at 6000x magnification and digital images were taken on a GATAN Orius camera. Representative cross-sectional planes of the sperm midpieces were taken for an n > 15 FOVs for each animal sample.

#### Sperm JC1 staining and flow cytometry

JC-1, a fluorescent, cationic, lipophilic dye, was used to demonstrate differences in mitochondrial membrane potential (MitoProbeTM JC-1 Assay Kit for Flow Cytometry, Invitrogen cat. number M34152). In brief, sperm collected fresh from swim-out were resuspended in 500 μM swim-out solution with final concentration of 3 μM JC-1 dye and gently rocked at 37°C for 20 minutes in the dark prior to two rounds of rinses (333 *× g* for 3 minutes). Per manufacturer instructions, CCCP (carbonyl cyanide m-chlorophenyl hydrazine) 25 μM final concentration was used as positive control to disrupt mitochondrial potential and to confirm that the JC-1 response was indeed sensitive. JC-1 exhibits potential-dependent accumulation of red fluorescent J-aggregates in mitochondria, indicated by a fluorescence emission shift from green (∼529 nm) to red (∼590 nm). In this way, mitochondrial depolarization can be inferred by a decrease in the red/green fluorescence intensity ratio, so sperm were analyzed using a BD Biosciences FACSCanto II and gated for green (FITC) and red (PE) fluorescence from a 488 laser. Percentage of sperm with defective mitochondrial membrane potential FITC-pos PE-neg were gated from a minimum of 200,000 events per replicate. Representative sperm from each gating strategy were also sorted directly onto microscope slides using BD Biosciences FACSAria II and imaged.

### Mammalian cell culture methods

All mammalian cells were cultured continuously at 37°C in a humidified incubator with 5% CO_2_. Immortalized fibroblasts were generated from mouse embryos isolated at E12.5 by sacrificing pregnant females and culturing the mouse embryonic fibroblasts (MEFs) as described previously (14, 95). Once established, these primary cells from wildtype, MRP9 single knockout and double knockout mice were immortalized by retroviral infection of MEFs with conditioned media from Ψ2-U195 cells producing the SV40 large T antigen. For overexpression in cell culture, plasmids for either *ABCC5* or *ABCC12* constructs were transfected into cell lines using PolyJet transfection reagent (SignaGen Laboratories, cat. number SL100688). Previously, human *ABCC5* ORF was cloned into pcDNA3.1(+)zeo and pEGFP-N1 plasmids for mammalian expression (14). Likewise, human *ABCC12* ORF was cloned into pcDNA3.1(+)zeo via EcorI and XbaI in a similar manner and subsequently epitope tagged with FLAG. Cells that were transfected with either vector or MRP expressing constructs could then be harvested up to 48 hours later for western blotting (protein expression), microscopy (localization) or other downstream assays. For attempting induction of endogenous expression of MRP5 or MRP9 in cell lines possibly capable of production (i.e. T-47D, A549, HCT116 and GC-2spd), cells were treated for 24 hours with 2 mM Sodium Butyrate (NaB) following methods from Bin Shi et al 2020 and then analyzed for expression and subcellular localization (72). For western blotting, cells were harvested in standard lysis buffer and followed standard immunoblotting procedure as described above. For polarization of MDCKII cells, trypsinized cells were seeded onto transwell cell culture inserts (Corning, COSTAR Polystyrene 6 well plates 24mm inserts, cat. number 3450) and allowed to grow to a monolayer in standard growth media (DMEM supplemented with 10% FBS) changed daily. Prior to reaching confluence, MDCKII cell’s monolayer integrity was determined daily by measuring the transepithelial electrical resistance (TEER) using an EVOM^2^ Epithelial Voltohmmeter device (World Precision Instruments, cat. number EVOM2). MDCKII cells were considered polarized and used for imaging when TEER values exceeded 200 Ω/cm^2^ resistance and were validated by checking for markers of polarization. Immunofluorescence was performed as follows: cells seeded onto coverslips (or transwells) were rinsed with ice cold DPBS (and subsequently in between each additional step) and immediately fixed in freshly prepared 4% PFA for greater than 30 minutes; cells were then quenched with 0.1 M ethanolamine for 5 minutes, twice; permeabilized with 0.1% Triton-X 100 in DPBS solution for 10 minutes; and blocked in blocking solution of 3% BSA:Superblock (SuperBlock Buffer in PBS, Thermo Scientific, cat. number 37515) for greater than one hour at room temperature or overnight at 4°C; incubated with primary antibody (rat monoclonal anti-MRP9 M9I-38, 1:10 in blocking solution) overnight at 4°C; secondary antibody (Alexa fluorophore-conjugated goat anti-rat, 1:2000 in blocking solution) for 1 hour at RT; and subsequently counterstained with DAPI (1:30,000 of 5 mg/ml) for 3 minutes and mounted using Antifade reagent (ProLong Gold Antifade Reagent, Invitrogen, cat. number P36934). Images were taken and processed using an Airyscan980 SR confocal microscope (Zeiss).

### Metabolomics methods

#### Global untargeted metabolomics of mouse testes

Tissue was first extracted either in aqueous or organic methods. For aqueous extractions, 10 volumes of methanol:H_2_O (3:1) was added to the tissue sample and homogenized with ceramic beads (Precellys®24-Dual, PeqLab). Then 200 μl of the homogenate was transferred to a new tube where it was then vortexed and centrifuged at 15,000 RPM for 10 min at 4°C. The supernatant was transferred to sample vial for ultra high performance liquid chromatography coupled with mass spectrometry (UHPLC-MS) analysis. Analytes were separated with either a CSH C18 column or a BEH amide column for more nonpolar and polar species, respectively. For the first aqueous method, separation was achieved using an ACQUITY UPLC CSH C18 Column (1.7 μm, 2.1 mm × 100 mm). Mobile phase A was water (contained 0.1% formic acid) and mobile phase B was acetonitrile (contained 0.1% formic acid). The gradient was: 0 to 1 min, 5% B; 1.1 to 10 min, 5% to 95 % B; 10.1 to 13 min, 95 % B; 13.1 to 13.5 min, 95 to 5% B; 13.6 to 15 min, hold at 5% B. The flow rate was 0.5 mL/min. The column was maintained at 50°C and the auto-sampler was kept at 10°C. A 5 μL injection was used for all samples. For the second method of aqueous analysis, the separation was achieved using an ACQUITY UPLC BEH amide Column (1.7 μm, 2.1 mm × 100 mm). Mobile phase A was water (contained 0.1% formic acid) and mobile phase B was acetonitrile (contained 0.1% formic acid). The gradient was: 0 to 1 min, 99% B; 1 to 10 min, 99% to 30 B; 10.1 to 12 min, hold at 99% B. The flow rate was 0.5 mL/min. The column was maintained at 45°C and the auto-sampler was kept at 10°C. A 5 μL injection was used for all samples. For both aqueous sample preps, data was acquired in both positive (HDMSe) and negative (HDMSe) mode. The capillary voltages were separated for positive (0.8 kV) and negative (0.8 kV) and sampling cone voltage was 40 V. Nitrogen at a flow of 800 L/h was used as the desolvation gas with a constant desolvation temperature of 500°C. The source temperature was set at 150°C. Data were acquired over the *m/z* range of 50-1000. For organic extraction (lipid) methods, likewise 10 volumes of methanol:H_2_O (3:1) was added to the tissue sample and homogenized with ceramic beads (Precellys®24-Dual, PeqLab). However, now after homogenization 400 ul was transferred to a new tube and 500 μL of ice-cold methyl-tert-butyl ether (MTBE) is added, prior to incubation at 650 RPM for 1 hour. Additionally, 500 μL of ice-cold water was then added and incubated at 650 RPM for 15 min before phase separation was completed by centrifugation for 8 min at 8,000 RPM at 4°C. The top (organic) layer was removed and dried at room temperature under nitrogen. The recovered lipids were reconstituted in 200 μL of isopropanol:acetonitrile:water (2:1:1, v/v/v) and were transferred to sample vial for UPLC-MS analysis. The separation was achieved using a CORTECS HILIC Column (2.7 μm, 2.1 mm × 100 mm). Mobile phase A was water/acetonitrile (5:95, v/v) with 10 mM ammonium acetate and mobile phase B water/acetonitrile (50:50, v/v) with 10 mM ammonium acetate. The gradient was ramped from 0.1% to 20% B in 10 min, ramped to 80% B in 3 min, ramped back down to 0.1% B and held for 3 min. The flow rate was 0.5 mL/min. The column was maintained at 30°C and the auto-sampler was kept at 10°C. A 5 μL injection was used for all samples. Like the aqueous phase, data was acquired in both positive (HDMSe) and negative (HDMSe) mode. The capillary voltages were separated for positive (2.8 kV) and negative (1.9 kV) and sampling cone voltage was 30 V. Nitrogen at a flow of 900 L/h was used as the desolvation gas with a constant desolvation temperature of 550°C. The source temperature was set at 120°C. Data were acquired over the *m/z* range of 100-1500. Positive and negative mode data were basally analyzed by Progenesis QI and putative metabolite identification was searched by using HMDB and LIPID MAPS databases (delta < 5 ppm). The output marker lists were then manually sorted and annotated for more sophisticated downstream analysis with MetaboAnalyst (96–98). Multivariate data analysis, volcano plots, Variable Importance in Projection (VIP) heat maps and mummichog algorithm analysis to putatively annotate metabolites were performed using MetaboAnalyst 4.0 (35).

#### Measurement of retinoids from reproductive tissues

Two separate analyses were performed to determine the retinoid profile in wild type and double knockout mice. Testes and seminal vesicles were assayed for retinoic acid isomers and retinol and total retinyl esters. Tissue samples were stored in −80°C until processed. Only glass containers, pipettes, and syringes were used to handle retinoids. Extraction of retinoids was performed under yellow lights using a two-step liquid-liquid extraction that has been described in detail previously using 4,4-dimethyl-RA as an internal standard for RA and retinyl acetate as an internal standard for retinol and total retinyl ester (99–102). Briefly, for the extraction of retinoids, tissue was homogenized in 1 mL of 0.9% NaCl (normal saline) and to each homogenate aliquot, 3 mL of 0.025 M KOH in ethanol was added to the homogenate followed by addition of 10 mL hexane to the aqueous ethanol phase. The samples were vortexed and centrifuged for 1 to 3 min at 1,000 rpm in a Dynac centrifuge (Becton Dickinson) to facilitate phase separation and pellet precipitated protein. The hexane (top) phase containing nonpolar retinoids [retinol and total retinyl esters (RE)] was removed. Then 4 M HCl (200 μL) was added to the remaining aqueous ethanol phase, samples were vortexed, and then polar retinoids (RA) were removed by extraction with a second 10 mL aliquot of hexane as described above. Organic hexane phases were evaporated under nitrogen while heating at approximately 30°C in a water bath (model N-EVAP 112, Organomation Associates, Berlin, MA, USA). All samples were resuspended in 60 μL acetonitrile. Levels of RA were determined by liquid chromatography-multistage tandem mass spectrometry (LC-MRM^3^) which is an LC-MS/MS method utilizing two distinct fragmentation events for enhanced selectivity (99). RA was measured using a Shimadzu Prominence UFLC XR liquid chromatography system (Shimadzu, Columbia, MD) coupled to an AB Sciex 6500+ QTRAP hybrid triple quadrupole mass spectrometer (AB Sciex, Framingham, MA) using atmospheric pressure chemical ionization (APCI) operated in positive ion mode as previously described (99). For the LC separation, the column temperature was controlled at 25°C, the autosampler was maintained at 10°C and the injection volume was typically 20 μL. All separations were performed using an Ascentis Express RP-Amide guard cartridge column (Supelco, 50 × 2.1 mm, 2.7 μm) coupled to an Ascentis Express RP-Amide analytical column (Supelco, 100 × 2.1 mm, 2.7 μm). Mobile phase A consisted of 0.1% formic acid in water, and mobile phase B consisted of 0.1% formic acid in acetonitrile. Endogenously occurring retinoid isomers including all-trans-retinoic acid (RA), 9-*cis* retinoic acid, 13-*cis* retinoic acid, and 9,13-di*-cis* retinoic acid are resolved using a gradient separation at a flow rate of 0.4 mL min^−1^ with gradient conditions described previously (99). The APCI source conditions and MRM^3^ detection parameters were as previously described where the MRM^3^ transition for RA was *m/z* 301.1 → *m/z* 205.1 → *m/z* 159.1 and for 4,4-dimethyl RA was *m/z* 329.2 → *m/z* 151.2 → *m/z* 100.0 (99).

Levels of retinol and total retinyl esters (RE) were quantified via HPLC-UV according to previously published methodology (102, 103). Note: total retinyl esters is 90+% retinyl palmitate. Retinol and RE were resolved by reverse-phase chromatography (Zorbax SB-C18, 4.6 × 100 mm, 3.5 μm) on a Waters Acquity UPLC H-class system and were quantified by UV absorbance at 325 nm. Analytes were separated at 1 mL min^−1^ with a gradient separation as described previously with a typical injection volume of 30 μL. The amount of retinoic acid (RA), retinol (ROL) and total retinyl ester (RE) was normalized per gram of tissue.

#### Targeted metabolomics of subcellular fractionation of testes

High-Throughput Targeted Metabolomics were performed on subcellular fractions isolated from WT and DKO testes. This subcellular fractionation was performed as described above with minor modifications, generating “Total Homogenate”, “740 *× g*”, “Membrane Prep”, “Mitochondria” and “MAMs” fractions. In order to maintain compatibility with downstream mass spectrometry analysis, each fraction was collected in ice cold 0.9% saline for its final centrifugation and subsequently was rinsed and diluted in ice cold 0.9% saline instead of lysis buffer prior to freezing. The concentration of metabolites and lipids were then determined with mass spectrometry using MxP® Quant 500 kit (Biocrates Life Sciences AG, Innsbruck, Austria). The kit enables quantification of approximately 630 endogenous metabolites belonging to 26 different biochemical classes: alkaloids (1), amine oxides (1), amino acids (20), amino acids related compounds (30), bile acids (14), biogenic amines (9), carbohydrates (1), carboxylic acids (7), cresols (1), fatty acids (12), hormones (4), indoles and derivatives (4), nucleobases and related compounds (2), vitamins (1), acylcarnitines (40), lysophosphatidylcholines (14), phosphatidylcholines (76), sphingomyelins (15), ceramides (28), dihydroceramides (8), hexosylceramides (19), dihexosylceramides (9), trihexosylceramides (6), cholesteryl esters (22), diglycerides (44), triglycerides (242). The method combines direct infusion coupled with tandem mass spectrometry (DI – MS/MS) and reverse phase liquid chromatography coupled with tandem mass spectrometry workflows (LC – MS/MS). Extraction was carried out per manufacturer’s instructions. Tissue samples were homogenized with ceramic beads (Precellys®24-Dual, Bertin Technologies SAS, France) prior to analysis. Briefly, mitochondrial fractions and tissue homogenates were thawed over ice and 30 μl of the sample was loaded onto the filter plate in 96 well format, impregnated with deuterated internal standards and dried under a stream of nitrogen in a positive pressure manifold. A 5% solution of phenylisothiocyanate in ethanol:water:pyridine (1:1:1, v:v:v) was added for derivatization of metabolites. Extraction was done with 5 mM ammonium acetate in methanol. The extracts were eluted with positive pressure manifold, followed by dilution steps for LC-MS/MS and DI-MS/MS acquisitions. The tandem mass spectrometry platform consisted of a Waters I Class Ultra performance liquid chromatography (UPLC) coupled to a Waters TQ-XS tandem quadrupole mass spectrometer (Waters Corporation, Milford, MA). MetIDQ software (Biocrates Life Sciences AG, Innsbruck, Austria) was used to register the samples, analyze the data and validate the assay. All reagents used were of analytical grade and mobile phases used were of LC-MS grade.

### Transcriptomics and associated methods

#### RNAseq of mouse testes

RNA was extracted from frozen tissues following standard Trizol chloroform protocol (Invitrogen, cat. number 15596026) and aqueous fractions were then taken directly into Qiagen RNA plus extraction kit (Qiagen, cat. number 74136) for maximum purity. Total RNA quality and quantity were tested on an Agilent Bioanalyzer 2100 System (Agilent cat. number G2939BA) using RNA Nano Chip On-Chip Electrophoresis trays for samples with RIN scores greater than 8 prior to processing for library prep/construction and sequencing. RNA-seq was performed at NISC using Illumina NovaSeq6000. FastQC, version 0.11.7, was used as an additional bioinformatics quality control on output reads from sequencing. The reads were then aligned to the mouse mm10 GRCh38 reference genome for gene annotation using STAR (Spliced Transcripts Alignment to a Reference), version 2.7.0f, and numbers of reads in transcripts per million (TPM) mapped to genes were counted generating matrixes for all samples. Differentially expressed genes were then identified via both manual export, sorting and fold change annotation, as well as processed via DESeq2, version 1.12.3, for more complex analysis. DESeq2 was run using R (version 3.6.1) and Bioconductor (version 3.4) with BioInstaller (version 1.24.0) for volcano plotting and statistical analysis of differential changes utilizing adjusted *P* value < 0.01 and False Discovery Rate (FDR) cutoffs of 0.05. The complete RNA sequencing results have been depositing to the NCBI Gene Expression Omnibus under accession under GSE176740.

#### Pathway analysis

Gene Ontology (GO) enrichment and network analysis of RNAseq results were performed using both Ingenuity Pathway Analysis (IPA) software package (Qiagen version 1-16) and manually developed excel pipeline workflows. For IPA analysis, statistically significant gene lists by tissue were manually filtered for minimum threshold of 10 TPM, sorted by *P* value and uploaded along with gene expression fold change for analysis. For pipeline pathway analysis, gene lists for GO terms and associated pathways of interest were downloaded from MGI (104–106), RGD (107) as well as manually added from literature review of relevant publications. These all-encompassing lists were then used as target gene IDs for filtered call look up functions from RNAseq gene expression RSEM matrixes. In brief, genes with matching IDs which passed threshold of expression in at least one sample condition and were statistically significant, were output in a heatmap with *P* value and relative fold change displayed.

#### Motif analysis

The MEME Suite and HOMER motif platforms were used to unbiasedly identify conserved motifs via discovery analysis of enriched sequences from differentially expressed genes. More specifically, we generated gene lists (either globally or mitochondria targeted) and divided them into upregulated and downregulated genes and extracted their associated sequence FASTA files. Then 1000bp upstream of ORF and 50bp downstream were used to run MEME or HOMER via Perl scripts on NIH High Performance Cluster (HPC) to discover the top 5 ungapped enriched putative motif sequences. These sequences were then further analyzed with the web-based suite tools TomTom and GoMo for MEME motif findings and manually curated for motif verification, which were further validated by checking and annotating output from the HOMER suite.

### Additional Bioinformatics and statistical analyses

All alignment and evolutionary analyses were conducted using MEGA × software version 10.2.4 (108–111). All data throughout are expressed as means ± SEM unless otherwise stated. Means of groups were assessed by using two-sided, unpaired student’s t-test or the equivalent analyses where applicable unless otherwise stated. A *P* value of < 0.05 when compared to baseline values or genotypes (WT) was considered statistically significant. All analyses were performed using Prism 9 software (GraphPad, version 9.1.0).

## Acknowledgments

We would like to acknowledge Ms. Lisa Garrett for conducting the pronuclear injections at NHGRI, NIH to generate the *Abcc12* mouse. We also thank Mr. Simon Beardsley for sharing the *C. elegans mrp-5* data. We acknowledge the Imaging Core Facility in the Department of Cell Biology and Molecular Genetics at the University of Maryland, College Park supported by Award Number 1S10OD025223-01A1 from the National Institutes of Health. This work was also done in part by utilizing the computational resources of the National Institutes of Health HPC Biowulf cluster (http://hpc.nih.gov). The MRP5 (M5I-10, M5-NC3) and MRP9 (M9I-27, M9I-38, M9II-3) antibodies used throughout were a generous gift from Piet Borst / Koen van de Wetering (Netherlands Cancer Institute). This work was supported by funding from the National Institutes of Health DK85035 and DK125740 (IH); HD077260 (MAK); the Utah Center for Iron and Heme Disorders was supported by funding from DK110858 (JP); and the Intramural Program of the National Human Genome Research Institute (DB). The funders had no role in study design, data collection and analysis, decision to publish, or preparation of the manuscript.

