## Supplemental Information for "MRP5 and MRP9 Play a Concerted Role in Male Reproduction and Mitochondrial Function"

**This PDF file includes:**

Figures S1 to S8
Tables S1 to S4
Legend for Movie S1

**Other supplementary materials for this manuscript include the following:**

Movie S1

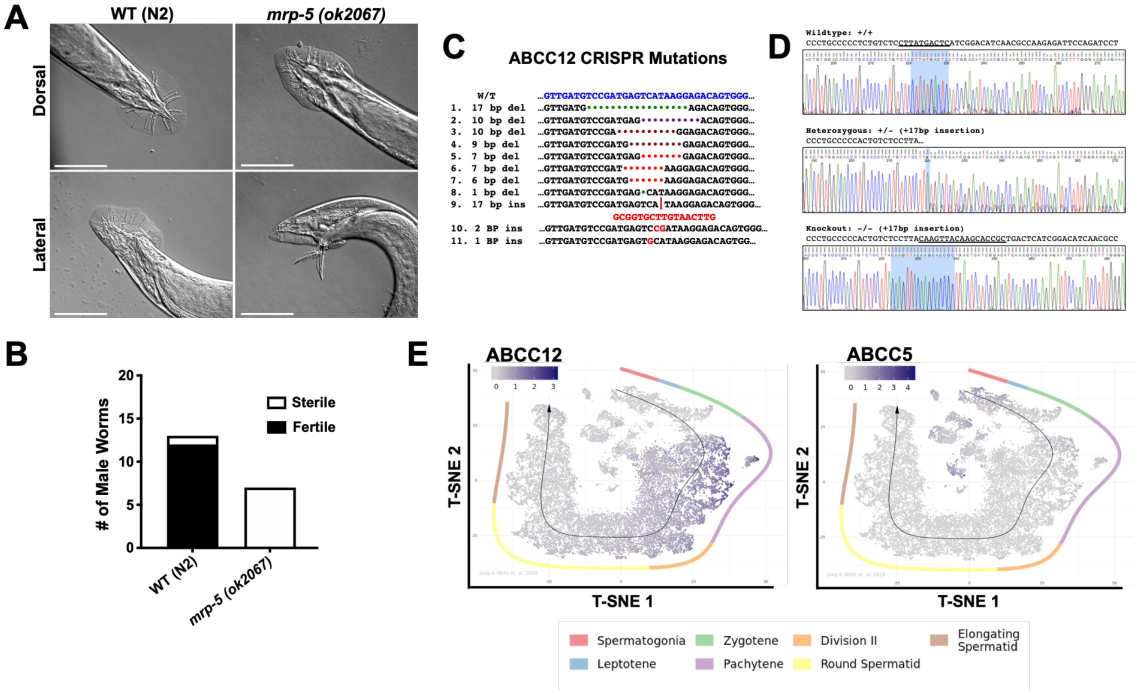


**Fig. S1. The generation of MRP9 knockout mice, a MRP5 paralog absent in worms**

**{A}** MRP-5 null worms (right) display reproduction related defects in the male tail with a reduced number of rays (top) and potentially defective spicule retraction after mating attempts (bottom) compared to WT (left). Scale bar equals 50 μm. **{B}** 7 Males of each indicated strain were picked onto a plate with one sperm exhausted WT hermaphrodite per replicate; wildtype n = 13 (12 / 1) *mrp-5* n = 7 (0 / 7). Presence of progeny was used as a measure of male mating success due to the fact that hermaphrodites were sperm exhausted. **{C}** DNA sequence of *Abcc12* target site and the sequences of the CRISPR induced mutations generated in the founder animal lines. **{D}** Sequencing chromatograms of RT-PCR from testes of *Abcc12* WT, HET and KO mice confirming mutation. **{E}** Single cell RNAseq testes atlas output of *Abcc12* (left) and *Abcc5* (right) gene expression profiles along the temporospatial axis of spermatid maturation taken from Jung et al 2019 dataset.

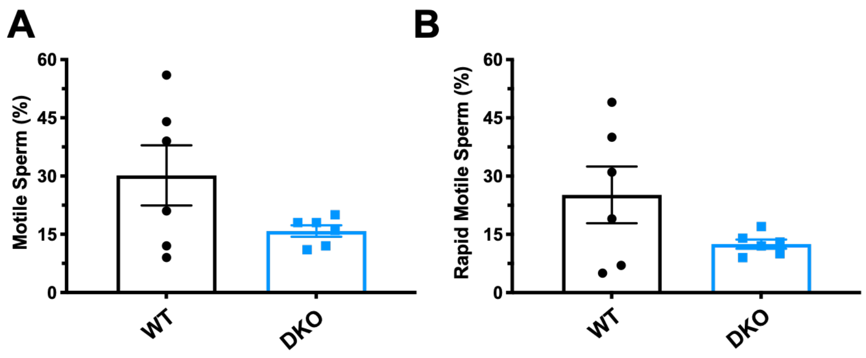


**Fig. S2. DKO sperm show trends in reduced motility**

**{A}** Percentage of "progressive motile" capacitated WT and DKO sperm assessed by IVOS computer assisted sperm analyzer system. **{B}** Percentage of "rapid motile" capacitated sperm assessed by IVOS computer assisted sperm analyzer system.

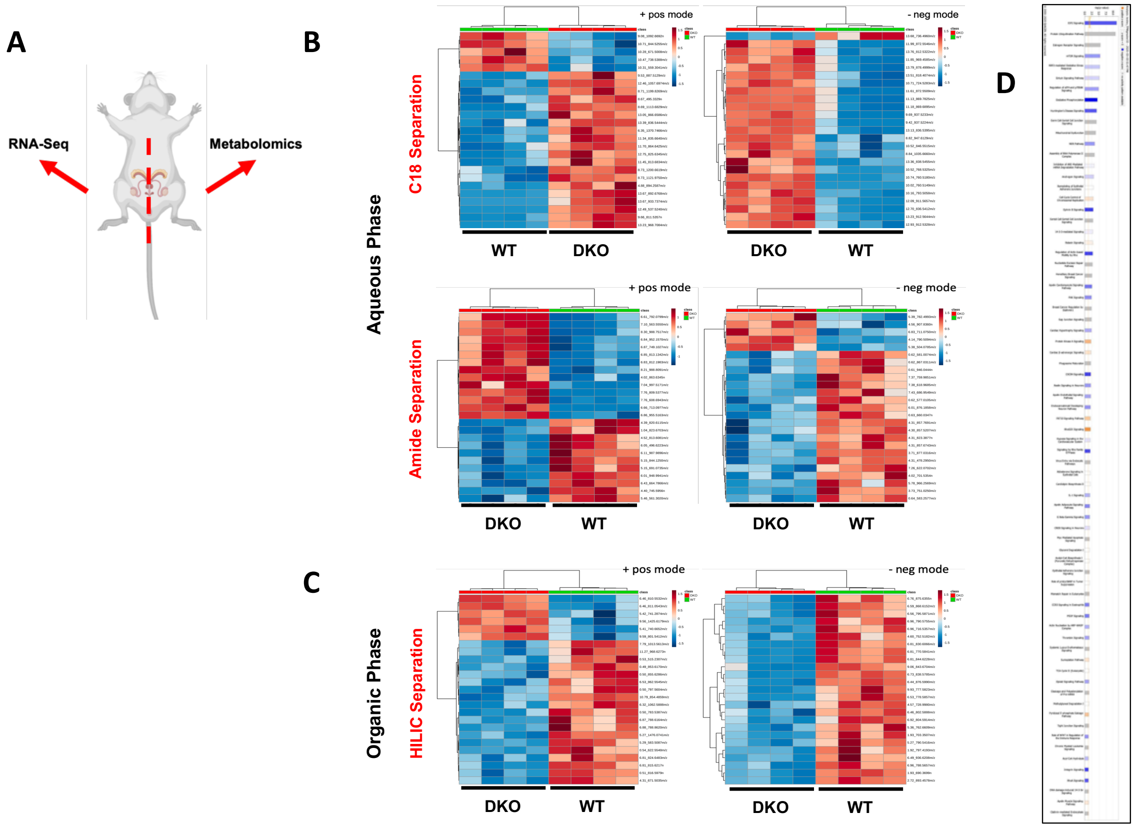


**Fig. S3. Metabolomics and RNAseq from the same animal highlight pervasive perturbations in the testes of DKO mice**

**{A}** Schematic of paired testes from same mice for both metabolomics and RNAseq analysis. **{B}** Variable Importance in Projection (VIP) heatmaps of differential metabolites from aqueous phase extractions separated on C18 (top) or amide (bottom) columns. Samples analyzed via mass spectrometry in positive ion (left) or negative ion (right) mode from testes in WT and DKO mice. **{C}** Variable Importance in Projection (VIP) Heatmaps of differential metabolites from organic phase extractions separated on a HILIC column. Samples analyzed via mass spectrometry in positive ion (left) or negative ion (right) mode from testes in WT and DKO mice. **{D}** All significantly altered GO pathways by Qiagen IPA analysis of testes RNAseq differentially expressed genes.

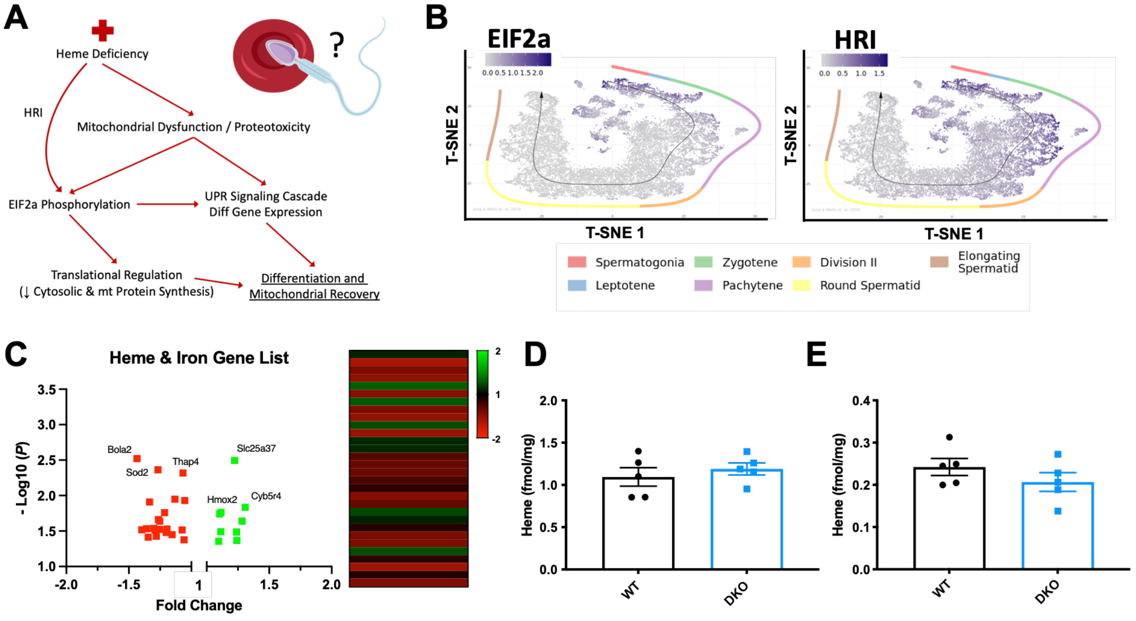


**Fig. S4. Probing heme / iron homeostasis as a culprit for EIF2 signaling in the testes**

**{A}** Schematic of canonical heme dependent EIF2a regulation of mitochondrial homeostasis and differentiation in the erythron lineage, which has not yet been examined in spermatozoa differentiation / proliferation. **{B}** Gene expression profiles for *Eif2a* (left) and *Eif2ak1* (right) along the temporospatial axis of spermatid maturation from single cell RNAseq of the testes taken from Jung et al 2019. **{C}** Output of pathway pipeline analysis investigating all “heme”, “heme binding” and “iron homeostasis” GO pathway gene lists. Gene lists were curated for integration with RNA expression data. Real expression was validated by thresholding a minimum 10 transcripts per million and statistically significant genes ($P < 0.05$) fold change were output in volcano plot and heatmap with 30 total genes identified, sorted by ascending $P$ value. **{D}** Heme content of whole testis from DKO and WT mice by oxalic acid quantification, n=5 animals per genotype. **{E}** Heme content of seminal vesicles from DKO and WT mice by oxalic acid quantification, n=5 animals per genotype.

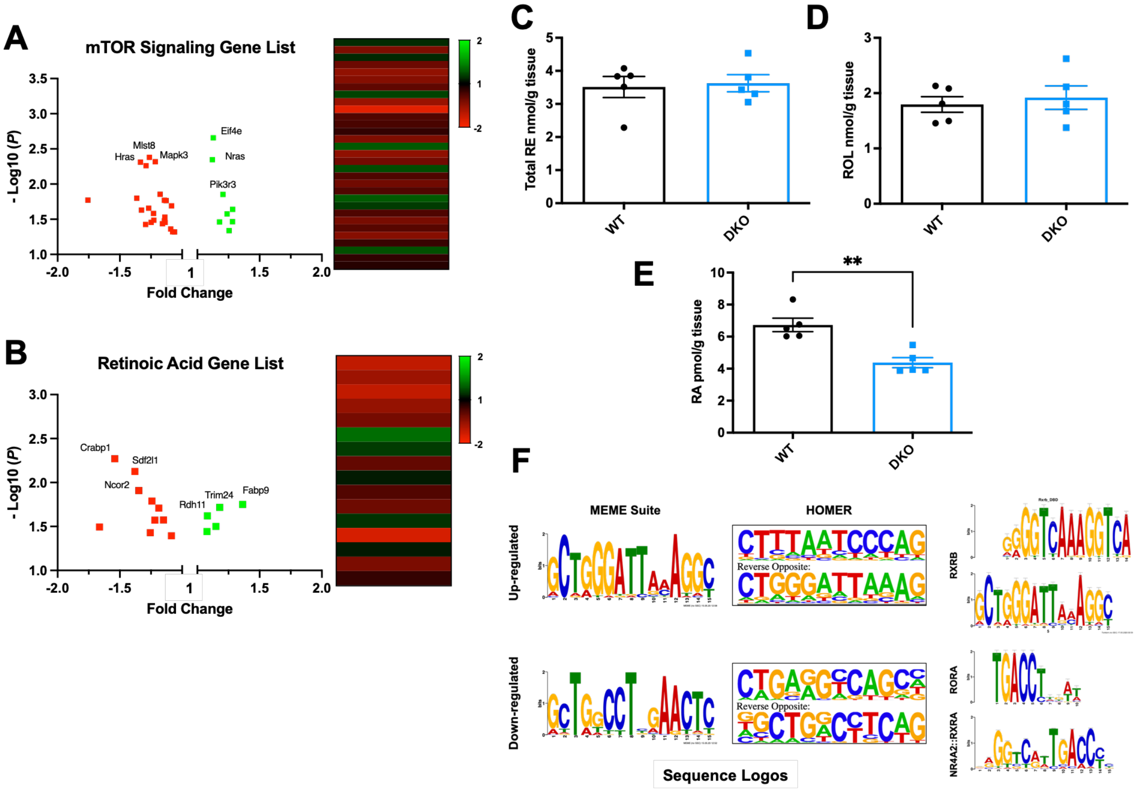


**Fig. S5. Retinoic acid homeostasis and signaling are altered in DKO testes**

**{A}** Schematic Output of pathway pipeline analysis investigating all "mTOR signaling" GO pathway gene lists. Gene lists were curated for integration with RNA expression data. Real expression was validated by thresholding a minimum 10 transcripts per million and statistically significant genes (*P* < 0.05) fold change were output in volcano plot and heatmap with 31 total genes identified, sorted by ascending *P* value. **{B}** Output of pathway pipeline analysis investigating all "Retinoic Acid", "Retinoid Metabolism" and "Retinoic Acid Signaling" GO pathway gene lists. Gene lists were curated for integration with RNA expression data. Real expression was validated by thresholding a minimum 10 transcripts per million and statistically significant genes (*P* < 0.05) fold change were output in volcano plot and heatmap with 15 total genes identified, sorted by ascending *P* value. **{C}** Quantification of total retinyl esters from WT and DKO testes, n=5 animals per genotype. **{D}** Quantification of Vitamin A (retinol) from WT and DKO testes, n=5 animals per genotype. **{E}** Quantification of all-*trans* retinoic acid from WT and DKO testes, n=5 mice, ** *P* value = 0.0025. **{F}** Upstream regulatory sequences (5` 1000bp) from all significantly up-regulated and down-regulated genes were processed to identify enriched un-gapped sequences conserved across transcripts with either MEME Suite (left) or HOMER (center). The top five statistically significant DNA sequences of each were then analyzed and aligned for querying against known binding motifs. We identified significant enrichment of retinoic acid related binding motifs including putative RXRs, and RORA binding sequences (right).

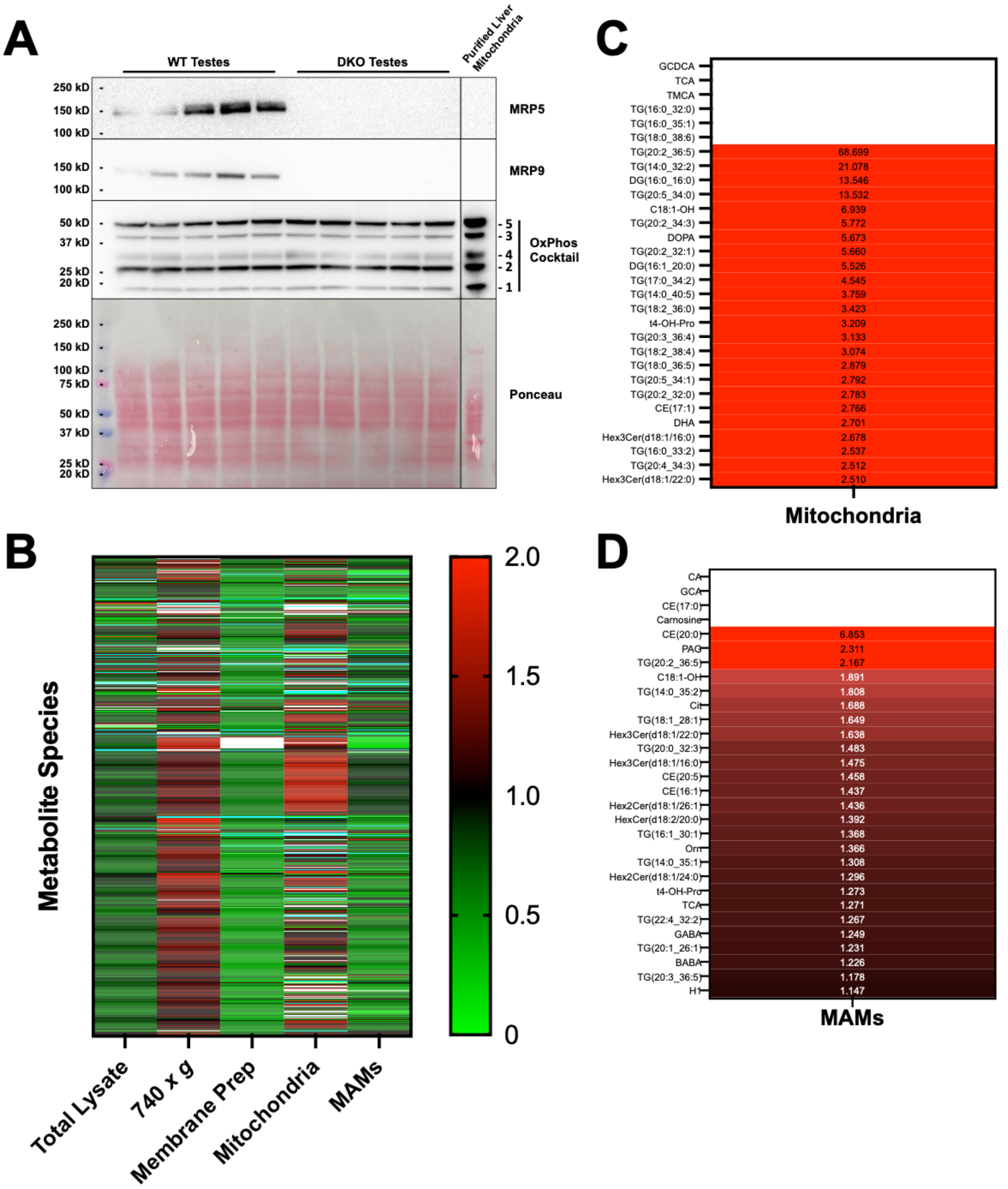


**Fig. S6. Target high-throughput metabolomics of testes reveals triglyceride accumulation in subcellular fractions**
**{A}** Immunoblotting of testes total lysates probing with anti-Total OxPhos Complex Kit, MRP5 and MRP9 antibodies. Complex Kit cocktail targets are premixed mouse monoclonal antibodies (#1 – Complex I, C-I-20 ND6; #2 – Complex II, C-II-30 FeS; #3 – Complex III, C-III-Core 2; #4 – Complex IV, C-IV-1; #5 – Complex V, C-V-a) and targets are labeled 1-5 on right hand side of the immunoblot. **{B}** Targeted high-throughput metabolomics of subcellular fractions from WT and DKO testes, utilizing Biocrates MxP® Quant 500 kit with quantification of 630 individual species, output represented as fold change normalized to total protein. **{C}** Top 30 accumulated metabolites and lipid species in the mitochondria of DKO testes quantified from subpanel B, fold change is annotated for each species while empty white bars represent infinity. **{D}** Top 30

accumulated metabolites and lipid species in the MAMs of DKO testes quantified from subpanel B, fold change is annotated for each species while empty white bars represent infinity.

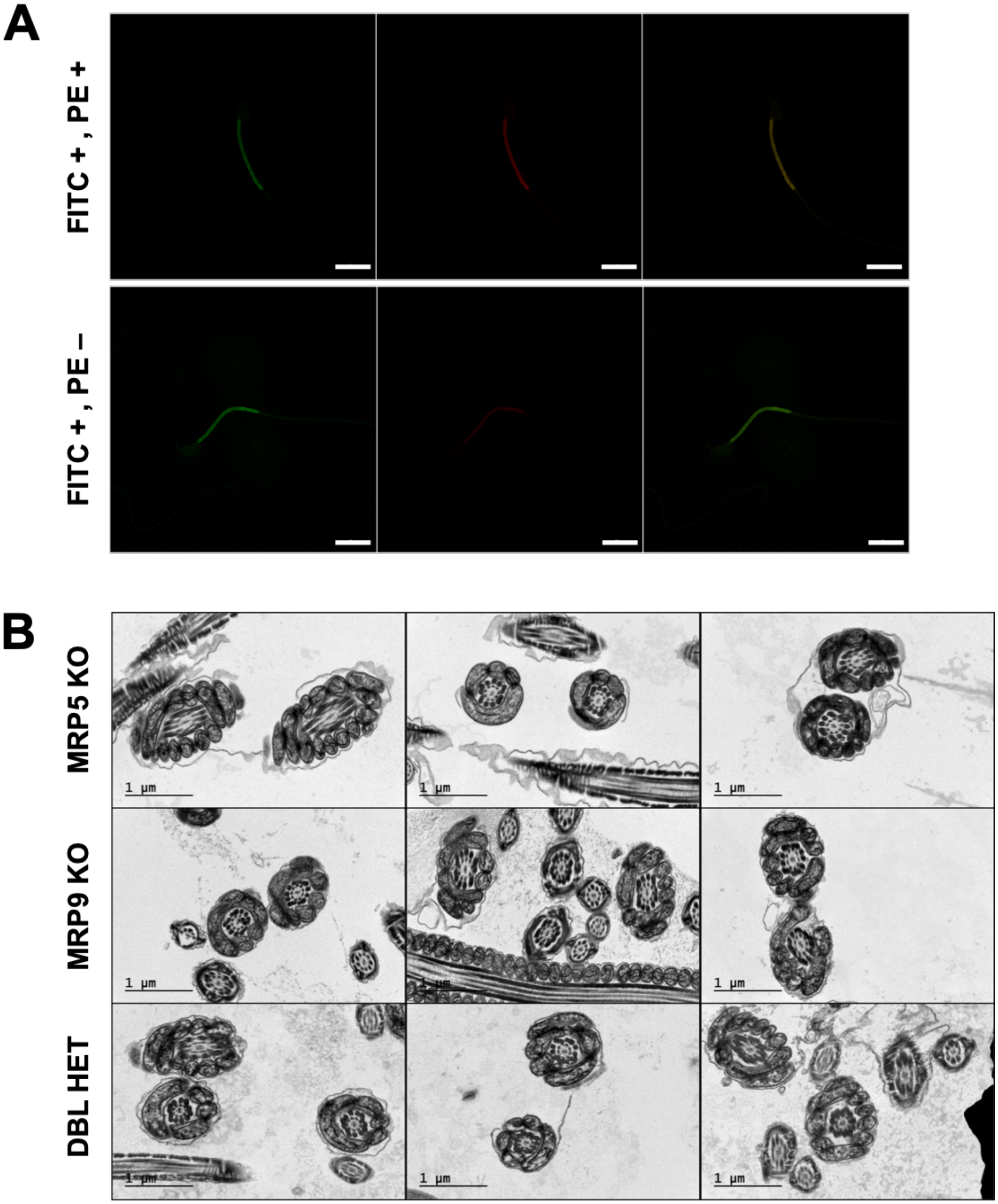


**Fig. S7. Additional sperm characterization via JC-1 staining and TEM imaging**
**{A}** Representative JC-1 staining of spermatozoa from **Figure 6A**, Q2 FITC + , PE + (top) and Q3 FITC + , PE – (bottom), scalebar equals 10 μm. **{B}** TEM of swum-out caudal epididymal spermatozoa from MRP5 KO (top), MRP9 KO (middle), and Double Het (bottom). Cross sections of the sperm midpiece visualize cristae and mitochondrial morphology of the mitochondrial sheath. Representative images from at least 15 FOVs per sample, scalebar equals 1 μm.

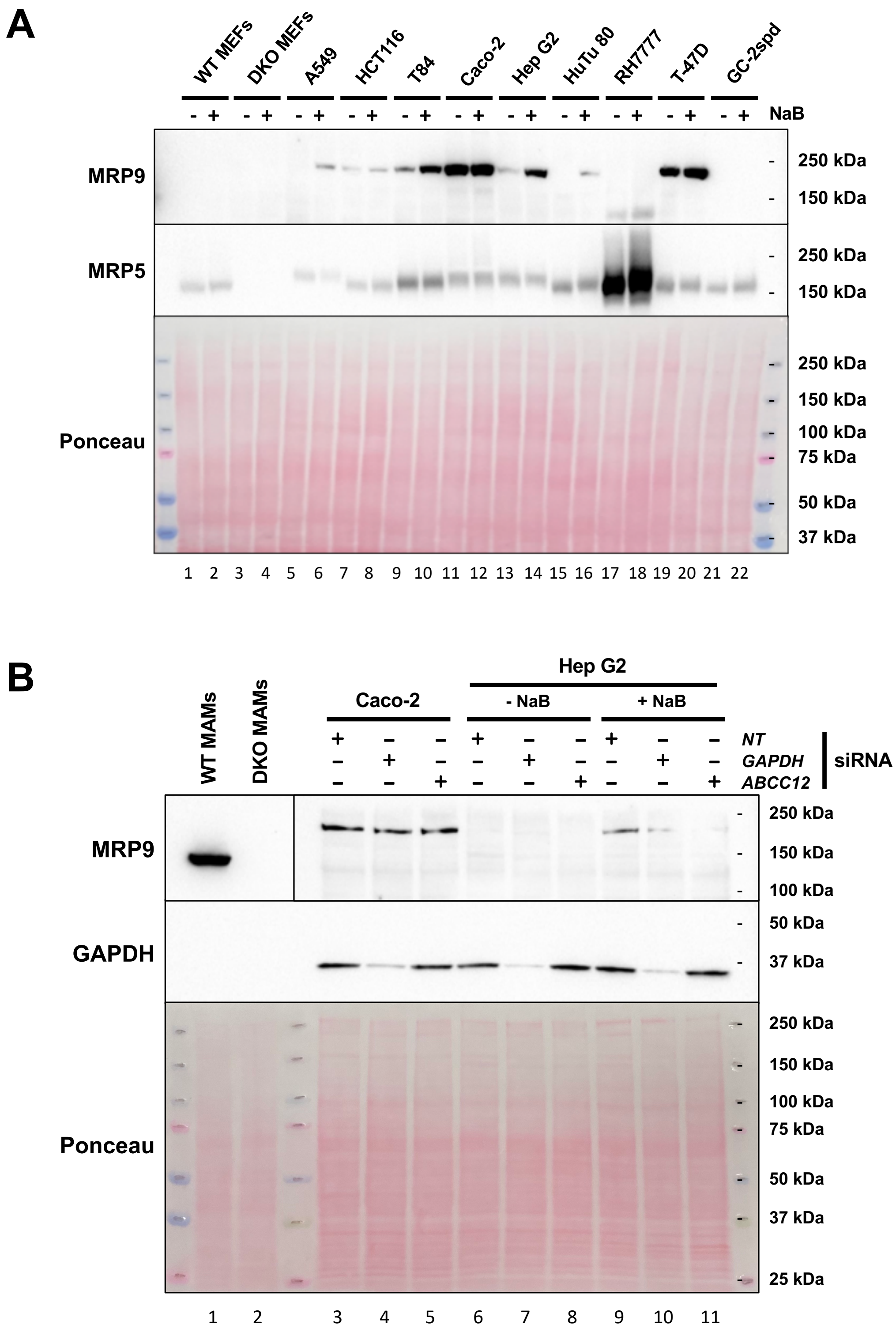


**Fig. S8. Endogenous expression of MRP9 cannot be detected or induced in cell culture** **{A}** Immunoblot of cell lines treated with either 2mM Sodium Butyrate (NaB) or PBS control for 24 hours; membrane was probed for MRP9 and MRP5. **{B}** Immunoblot analysis of selected cell lines with or without 4mM NaB and treated with Non-Targeted (NT), *GAPDH* or *ABCC12* siRNA for 48hours; membrane was probed for MRP9 and GAPDH protein levels.

**Table S1. Identifiable metabolites from MetaboAnalyst mummichog processing**

| Extraction Method | Compound KEGG ID | Matched form | Name / ID | pValue | Fold Change |
|---|---|---|---|---|---|
| Aqueous Pos | G00526 | M+NaCl[1+] | **Glycan - (GlcA)2 (GlcNAc)2** | 0.0006 | 3.3345 |
| Amide Pos | C11039 | M+CH3COO[-] | **2'-Deoxy-5-hydroxymethylcytidine-5'-triphosphate** | 0.0009 | -2.1258 |
| Amide Pos | C15976 | M+Cl[-] | **2-Methyl-1-hydroxypropyl-TPP** | 0.0017 | 2.7099 |
| Amide Pos | C11039 | M-H2O-H[-] | **2'-Deoxy-5-hydroxymethylcytidine-5'-triphosphate** | 0.0022 | -2.0490 |
| Amide Pos | C00127 | M+Br[-] | **Oxidized glutathione** | 0.0034 | -2.4940 |
| Amide Pos | G00044 | M+CH3COO[-] | **IV2-a-Fuc-Lc4Cer; Type IH glycolipid** | 0.0057 | -2.8923 |
| Amide Pos | G00055 | M+CH3COO[-] | **IV2-a-Fuc-nLc4Cer; Type IIH glycolipid** | 0.0057 | -2.8923 |
| Amide Pos | G00060 | M+CH3COO[-] | **III3-a-Fuc-nLc4Cer; Lacto-N-fucopentaosyl III ceramide** | 0.0057 | -2.8923 |
| Amide Pos | G10770 | M-2H[2-] | **(GlcNAc)3 (LFuc)1 (Man)3 (Asn)1** | 0.0060 | -2.3658 |
| Amide Pos | C05261 | M(S34)-H[-] | **3-Oxotetradecanoyl-CoA** | 0.0061 | -2.9834 |
| Amide Pos | C05261 | M(Cl37)-H[-] | **3-Oxotetradecanoyl-CoA** | 0.0061 | -2.9834 |
| Amide Pos | C04646 | M-H+O[-] | **Thioinosinic acid** | 0.0064 | -2.1401 |
| Amide Pos | C16618 | M-H[-] | **6-Thioxanthine 5'-monophosphate** | 0.0064 | -2.1401 |
| Amide Pos | C00387 | M+Br[-] | **Guanosine** | 0.0064 | -2.0488 |
| Aqueous Pos | C05504 | M+NaCl[1+] | **Estriol-16-Glucuronide** | 0.0070 | -2.6085 |
| Aqueous Pos | C11376 | M-HCOOH+H[1+] | **SN38 glucuronide** | 0.0070 | -2.6085 |
| Aqueous Pos | C16327 | M(C13)+2H[2+] | **OPC8-CoA** | 0.0070 | -2.6085 |
| Amide Pos | C00655 | M+Br[-] | **Xanthylic acid** | 0.0078 | 8.4958 |
| Amide Pos | C21750 | M+Cl37[-] | **5-Fluorodeoxyuridine diphosphate** | 0.0078 | 8.4958 |
| Amide Pos | C14855 | M+Br81[-] | **4,5-Dihydro-4-hydroxy-5-S-glutathionyl-benzo[a]pyrene** | 0.0080 | 7.1469 |
| Amide Pos | C14856 | M+Br81[-] | **7,8-Dihydro-7-hydroxy-8-S-glutathionyl-benzo[a]pyrene** | 0.0080 | 7.1469 |
| Aqueous Pos | G01945 | M[1+] | **(Gal)2 (GlcNAc)2 (S)3** | 0.0082 | 3.8200 |
| Amide Pos | G00159 | M-H2O-H[-] | **(Gal)2 (GalNAc)1 (GlcA)2 (Xyl)1 (Ser)1** | 0.0085 | -3.2323 |
| Amide Pos | G00163 | M-H2O-H[-] | **(Gal)2 (GlcA)2 (GlcNAc)1 (Xyl)1 (Ser)1** | 0.0085 | -3.2323 |
| Amide Pos | C11174 | M-H2O-H[-] | **1-Diphosinositol pentakisphosphate** | 0.0086 | -2.5534 |
| Amide Pos | C11526 | M-H2O-H[-] | **5-Diphosphoinositol pentakisphosphate** | 0.0086 | -2.5534 |
| Amide Pos | C00091 | M(C13)-H[-] | **Succinyl-CoA** | 0.0089 | 16.1675 |
| Amide Pos | C00683 | M(C13)-H[-] | **Methylmalonyl-CoA** | 0.0089 | 16.1675 |
| Amide Pos | C01213 | M(C13)-H[-] | **L-methylmalonyl-CoA** | 0.0089 | 16.1675 |
| Amide Pos | C03691 | M+Cl37[-] | **CMP-N-glycoloylneuraminate** | 0.0096 | -3.3747 |
| Amide Pos | C00877 | M-H[-] | **Crotonoyl-CoA** | 0.0113 | -2.3923 |
| Amide Pos | C01144 | M-H2O-H[-] | **(S)-3-Hydroxybutyryl-CoA** | 0.0113 | -2.3923 |
| Amide Pos | C03460 | M-H[-] | **Methacrylyl-CoA** | 0.0113 | -2.3923 |
| Amide Pos | C06000 | M-H2O-H[-] | **(S)-3-Hydroxyisobutyryl-CoA** | 0.0113 | -2.3923 |
| Amide Pos | G00005 | M(S34)-H[-] | **(GlcNAc)2 (Man)3 (PP-Dol)1** | 0.0116 | 3.6595 |
| Amide Pos | G00005 | M(Cl37)-H[-] | **(GlcNAc)2 (Man)3 (PP-Dol)1** | 0.0116 | 3.6595 |
| Amide Pos | G00066 | M(S34)-H[-] | **(Gal)2 (Glc)1 (GlcNAc)2 (Cer)1** | 0.0116 | 3.6595 |
| Amide Pos | G00066 | M(Cl37)-H[-] | **(Gal)2 (Glc)1 (GlcNAc)2 (Cer)1** | 0.0116 | 3.6595 |
| Amide Pos | G00095 | M(S34)-H[-] | **IV3GalNAca-Gb4Cer** | 0.0116 | 3.6595 |
| Amide Pos | G00095 | M(Cl37)-H[-] | **IV3GalNAca-Gb4Cer** | 0.0116 | 3.6595 |
| Amide Pos | G00889 | M(S34)-H[-] | **(Gal)2 (GalNAc)1 (Glc)1 (GlcNAc)1 (Cer)1** | 0.0116 | 3.6595 |
| Amide Pos | G00889 | M(Cl37)-H[-] | **(Gal)2 (GalNAc)1 (Glc)1 (GlcNAc)1 (Cer)1** | 0.0116 | 3.6595 |
| Amide Pos | G02977 | M(S34)-H[-] | **(Gal)2 (GalNAc)2 (Glc)1 (Cer)1** | 0.0116 | 3.6595 |
| Amide Pos | G02977 | M(Cl37)-H[-] | **(Gal)2 (GalNAc)2 (Glc)1 (Cer)1** | 0.0116 | 3.6595 |
| Amide Pos | C05264 | M+CH3COO[-] | **(S)-Hydroxydecanoyl-CoA** | 0.0117 | 3.1082 |
| Amide Pos | C00327 | M+K-2H[-] | **Citrulline** | 0.0118 | 3.1441 |
| Amide Pos | C05266 | M-H+O[-] | **(S)-Hydroxyoctanoyl-CoA** | 0.0127 | -3.7376 |
| Aqueous Neg | G00149 | M+Br81[-] | **(GlcN)1 (Ino(acyl)-P)1 (Man)3 (EtN)1 (P)1** | 0.0144 | 3.1562 |
| Amide Pos | C01832 | M+HCOO[-] | **Lauroyl-CoA** | 0.0172 | 3.5276 |
| Aqueous Pos | C18043 | M(S34)+H[1+] | **Cholesterol sulfate** | 0.0177 | -3.0325 |
| Aqueous Pos | C18043 | M(Cl37)+H[1+] | **Cholesterol sulfate** | 0.0177 | -3.0325 |
| Amide Pos | C00183 | M+Na-2H[-] | **L-Valine** | 0.0224 | -2.1317 |
| Amide Pos | C00719 | M+Na-2H[-] | **Betaine** | 0.0224 | -2.1317 |
| Amide Pos | G00026 | M+Cl[-] | **(Gal)1 (GalNAc)1 (Neu5Ac)1 (Ser/Thr)1** | 0.0228 | -8.3344 |
| Aqueous Pos | G00158 | M+HCOONa[1+] | **(Gal)2 (GalNAc)1 (GlcA)1 (Xyl)1 (Ser)1** | 0.0235 | -2.4856 |
| Aqueous Pos | G00162 | M+HCOONa[1+] | **(Gal)2 (GlcA)1 (GlcNAc)1 (Xyl)1 (Ser)1** | 0.0235 | -2.4856 |
| Aqueous Neg | G00157 | M+Br[-] | **(Gal)2 (GlcA)1 (Xyl)1 (Ser)1** | 0.0237 | -2.2724 |
| Organic Pos | C16338 | M+HCOOK[1+] | **3-Oxo-OPC4-CoA** | 0.0251 | 1.8715 |
| Amide Pos | C04646 | M-H+O[-] | **Thioinosinic acid** | 0.0292 | -2.7960 |
| Amide Pos | C16618 | M-H[-] | **6-Thioxanthine 5'-monophosphate** | 0.0292 | -2.7960 |
| Organic Pos | G13036 | M[1+] | **(GlcA)2 (GlcN)1 (GlcNAc)1 (S)3** | 0.0423 | 2.4015 |
| Amide Pos | C02843 | M(C13)-H[-] | **Long-chain acyl-CoA** | 0.0429 | -1.9489 |
| Aqueous Neg | C05791 | M+Cl[-] | **D-Urobilinogen** | 0.0454 | -2.2582 |
| Amide Pos | C00942 | M(Cl37)-H[-] | **Cyclic GMP** | 0.0455 | 2.0594 |
| Aqueous Pos | G00063 | M[1+] | **IV3NeuAc,III3Fuc-nLc4Cer** | 0.0480 | 1.9138 |

Fold change and *P* values for statistically significant species identifiable to KEGG IDs based on mummichog analysis from all six untargeted metabolomics datasets of the testes. Peak intensity tables were uploaded to MetaboAnalyst suite and queried with a 5ppm mass tolerance for ID.

**Table S2. All significant pathways identified by integrative MetaboAnalyst analysis of metabolomics and RNAseq**

| MetaboAnalyst Testes Pathways | Total | Expected | Raw p | -LOG(p) | FDR | Impact |
|---|---|---|---|---|---|---|
| **Phosphatidylinositol signaling system** | 74 | 5.1195 | 0.0013121 | 6.6362 | 0.082863 | 0.57534 |
| **Drug metabolism - other enzymes** | 69 | 4.7736 | 0.0022033 | 6.1178 | 0.082863 | 0.30882 |
| **Pyruvate metabolism** | 45 | 3.1132 | 0.0029594 | 5.8228 | 0.082863 | 0.70455 |
| **Citrate cycle (TCA cycle)** | 42 | 2.9057 | 0.0068637 | 4.9815 | 0.13233 | 1.1463 |
| **Glycolysis or Gluconeogenesis** | 61 | 4.2201 | 0.0078766 | 4.8439 | 0.13233 | 0.65 |
| **Purine metabolism** | 169 | 11.692 | 0.039654 | 3.2276 | 0.37186 | 0.65476 |
| **Propanoate metabolism** | 48 | 3.3208 | 0.044216 | 3.1187 | 0.37186 | 0.59574 |
| **Inositol phosphate metabolism** | 69 | 4.7736 | 0.045331 | 3.0938 | 0.37186 | 0.27941 |
| **Drug metabolism - cytochrome P450** | 39 | 2.6981 | 0.048529 | 3.0256 | 0.37186 | 0.15789 |
| **Ether lipid metabolism** | 39 | 2.6981 | 0.048529 | 3.0256 | 0.37186 | 0.31579 |
| **Lysine degradation** | 49 | 3.3899 | 0.048696 | 3.0222 | 0.37186 | 0.3125 |

Output of all statistically significant pathways determined by MetaboAnalyst combined analysis of RNAseq gene lists with differential expression changes and metabolomics mummichog putative KEGG IDs.

**Table S3. Guide RNA sequences for targeting mouse MRP9 (*Abcc12*)**

| | |
|---|---|
| GENE NAME: **Abcc12**<br>ACCESSION: **NM_172912** | |
| MRP9_mouse sgRNA#1: | **5'CCAGCATCATCCACAGGATT 3'** |
| MRP9_mouse sgRNA#2: | **5'TTGATGTCCGATGAGTCATA 3'** |

**Table S4. List of primers used for genotyping mice**

| Primer | Sequence |
|---|---|
| mMRP5 FWD | CTAGAGTCTAATCCGTATTGG |
| mMRP5 REV | CCCGCAAATACATTCAAACC |
| mMRP5-HYG FWD | GCTTTCAGCTTCGATGTAGG |
| mMRP5-HYG REV | CGTCAGGACATTGTTGGAGC |
| mMRP9 FWD | GGTCAGCAGCTCCTGTAG |
| mMRP9 REV | CTTCCTCCAGGACCCTGA |

**Movie S1 (separate file). Immunofluorescence of cells transfected with MRP9 confirms novel subcellular localization in close proximity to mitochondria**

Movie of 3D rendering and orthogonal slices from Airyscan super resolution microscopy of HeLa cells transfected with human *ABCC12.* Mitochondria were stained with Mitotracker Deep Red FM immediately prior to fixation and immunofluorescence. Antibody probing for MRP9 and Calnexin was followed by Alexa-488 and Alexa-568 secondary antibodies respectively followed by DAPI counter staining prior to mounting.
